# Large-scale genome analyses provide insights into Hymenoptera evolution

**DOI:** 10.1101/2024.07.01.601370

**Authors:** Chun He, Yi Yang, Xianxin Zhao, Junjie Li, Yuting Cai, Lijia Peng, Yuanyuan Liu, Shijiao Xiong, Yang Mei, Zhichao Yan, Jiale Wang, Shan Xiao, Ziwen Teng, Xueke Gao, Hui Xue, Qi Fang, Gongyin Ye, Xinhai Ye

**Author notes:** Correspondence (X.H. Ye).

## Abstract

The order Hymenoptera includes a large number of species with diverse lifestyles and is known for its significant contributions to natural ecosystems. To better understand the evolution of this diverse order, we performed large-scale comparative genomics on 131 species from 13 superfamilies, covering most representative groups. We used these genomes to reveal an overall pattern of genomic change in terms of gene content and evolutionary rate throughout hymenopteran history. We identified genes that possibly contributed to the evolution of several key innovations, such as parasitoidism, wasp-waist, sting, and secondary phytophagy. We also discovered the distinct genomic trajectories between the clade containing major parasitoid wasps (Parasitoida) and stinging species (Aculeata) since their divergence, which are involved in many aspects of genomic change, such as rapidly evolving gene families, gene gain and loss, and metabolic pathway evolution. In addition, we explored the genomic features accompanying the three independent evolution of secondary phytophagy. Our work provides insights for understanding genome evolution and the genomic basis of diversification in Hymenoptera.

## Introduction

Hymenoptera is a diverse order of insects, which includes more than 153,000 described species, such as sawflies, wasps, ants, and bees^1–3^. These hymenopterans play a pivotal role in natural ecosystems, serving a variety of characters (e.g., herbivores, parasitoids, predators, and pollinators)^4,5^. The remarkable diversity of species and the apparent changes in lifestyles have prompted interest in the evolution of Hymenoptera. The current phylogenetic analyses based on various datasets have largely established the phylogenetic framework of Hymenoptera and identified several key innovative events in their evolution^3,4,6,7^. In brief, the origin of the Hymenoptera is estimated to be approximately 281 million years ago (Mya) in the Permian period^3^. It is likely that the ancestral hymenopterans are plant-feeders, and this lifestyle is still retained in the majority of sawflies, which represent the early branches of the Hymenoptera tree^3,4^. The emergence of parasitoidism occurred in the last common ancestor of Orussoidea and wasp-waisted wasps (also known as Apocrita)^3,4^. This transition from phytophagy to parasitoidism is regarded as a key innovation in the evolution of Hymenoptera and may have been a significant driver of species diversification, as approximately 70% of all described hymenopterans are parasitoids^3,4,8^. In Apocrita, species have evolved a highly constricted “waist” between the first and second abdominal segments, and this wasp-waist allows for flexibility in female ovipositors, which may be an important adaptation for parasitoid lifestyles^4,9^. Another morphological innovation is the acquisition of the venomous stinger (the modified female ovipositor from an egg-laying apparatus to a stinging apparatus) in Aculeata^3,4^. And some stinging species in the Aculeata (e.g., ants and certain groups of bees and wasps) have also evolved into eusociality independently^3^. Furthermore, it is important to note that following the evolution of the basal hymenopteran diet from phytophagy to parasitoidism (equivalent to carnivory), certain gall wasps, fig wasps, and pollen-collecting bees have independently evolved again to phytophagy (i.e., secondary phytophagy)^4^. A recent phylogenetic study has indicated that the evolution of secondary phytophagy had a significant influence on the diversification rate in Hymenoptera^4^. Overall, the evolution of Hymenoptera has been accompanied by several key innovations and evolutionary changes of lifestyles; however, the genetic mechanisms underlying these evolutionary processes remain largely unknown.

The sequencing of numerous hymenopteran genomes has opened new avenues for a deeper understanding of Hymenoptera evolution, yet a comprehensive, large-scale comparative genomic analysis of Hymenoptera is lacking. To fill this gap, we performed a large-scale comparative genomics study containing 131 species, which represent major lineages of Hymenoptera. Using the genomic data, we systematically investigated: (1) the general patterns of genome evolution in terms of gene content and evolutionary rate throughout hymenopteran history; (2) genomic changes occurred at the evolutionary branches associated with key innovations; (3) differences in genome evolution between two major clades in the Hymenoptera phylogeny, the Parasitoida and Aculeata; and (4) genomic features related to three independently evolutionary events of secondary phytophagy. Our findings not only offer a comprehensive picture of the genome evolution of hymenopteran insects but also highlight several genomic changes potentially related to evolutionary innovations, shedding light on the genomic basis of adaptation and diversification within this remarkable order.

## Results and Discussion

### Hymenoptera phylogenomics

To explore the evolutionary history of Hymenoptera, we obtained high-quality genomes for 131 hymenopteran insects from 13 superfamilies (29 families), covering many representative lineages with unique lifestyles such as sawflies, wasps, ants, and bees (Supplementary Table 1). These genomes were all publicly available, and 78 of them were sequenced using long-read sequencing technologies, showing the overall high contiguity (mean Contig N50 4.19 Mb). Furthermore, BUSCO (Benchmarking Universal Single-Copy Orthologs) assessments indicated that these genomes have a high level of gene completeness (highest 99.80%, lowest 85.10%, mean 97.26%, s.d. = 0.03) (Supplementary Table 1). In order to minimize the impact of potential gene annotation issues on subsequent analysis, we then employed an alignment-based approach to identify potential broken and chimeric genes^10^, correcting about 3.6% of total genes (Supplementary Table 2). After gene corrections, orthology inference produced 33,405 orthogroups (OGs, also referred to as gene families), with 25.1% of them present in over 70% of hymenopterans (Fig. 1, Supplementary Fig. 1, and Supplementary Table 3). Of the total number of genes analyzed, 78,789 (4.77%) were identified as singletons, meaning that no homolog was predicted in any genome.

**Fig. 1.**
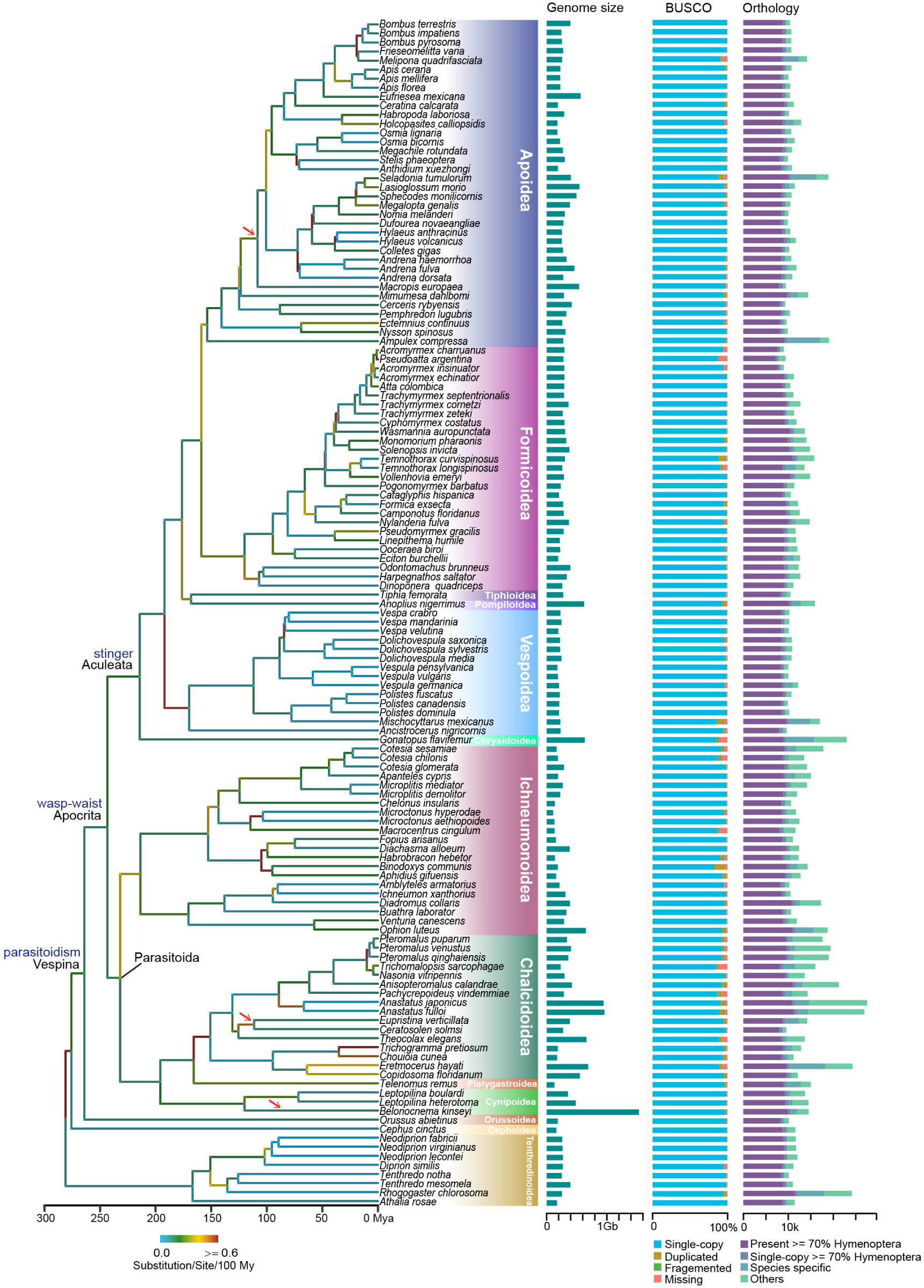
Phylogenetic analysis and genomic comparisons of Hymenoptera. The maximum-likelihood phylogenetic tree on the left was constructed with 1000 ultrafast bootstraps under a JTT+F+R10 model using protein sequences from 1002 single-copy genes. Almost all the nodes received 100% bootstrap support, except for nine. The branch color corresponds to the rate of substitutions per site per 100 million years along that particular branch. The divergence times among species were estimated using 13 calibration points. Several key innovations, such as parasitoidism, wasp-waist, and stinger, are marked on the phylogeny, and three independent transitions to secondary phytophagy are indicated by red arrows. On the right side, the figure shows i) genome assembly sizes of hymenopteran species; ii) BUSCO profiles for these genomes; and iii) orthology profiles. In the orthology analysis, the protein-coding genes of each species were subdivided to represent different types of orthology clusters.

We next reconstructed the phylogeny of hymenopterans based on protein sequences from 1002 single-copy genes (Fig. 1). Only nine (6.98%) nodes in this tree fail to receive 100% bootstrap support (Supplementary Fig. 2). Our species tree was mostly consistent with previous hymenopteran phylogenies using transcriptomic and mitochondrial data^3,7^. We showed that: (i) phytophagous sawflies from Tenthredinoidea and Cephoidea are placed in the early branches of Hymenoptera; (ii) Parasitoid Orussoidea is a sister to all wasp-waisted wasps, also known as Apocrita; and (iii) in Apocrita, parasitoid wasps from several superfamilies such as Ichneumonoidea, Chalcidoidea, Cynipoidea, and Platygastroidea form one clade (i.e., Parasitoida), and stinging species of Aculeata form its sister clade (Fig. 1). This phylogenetic relationship within Apocrita differs from the results of studies using ultraconserved elements^4,6^. These studies support that Ichneumonoidea is a sister to the clade containing all other species of Apocrita. There may be several reasons for this conflict, such as the type of data used and the samples analyzed. Further studies should expand the sampling to establish a more robust tree for Hymenoptera. In our subsequent analyses, we used the phylogenetic tree constructed from the genomic data, and the divergence times were estimated using 13 calibration points based on a previous study^3^ (Fig. 1 and Supplementary Fig. 2).

### Genomic changes throughout Hymenoptera history

We next investigated the changes in three different genomic features of protein-coding genes, including gene family evolution, protein domain rearrangement, and the evolutionary rate of protein-coding genes, to explore the overall dynamics of Hymenoptera genome evolution. First, we inferred the history of gene family evolution and identified 63,907 gene family expansions and 106,004 contractions (Supplementary Figs. 3–4). This pattern suggests a generally reductive mode of gene family evolution in Hymenoptera, which is also supported by a small-scale comparative genomic study of this order^11^. The pincer wasp *Gonatopus flavifemur* (Chrysidoidea, Dryinidae) is notable among hymenopterans for having the largest number of gene family expansion (736) and contraction (2564) events (Supplementary Fig. 3). We also estimated rates for gene family size evolution and found the average global rate of gene gain and loss of gene families in Hymenoptera was estimated to be 58.29 gene gain and loss per million years (My). The rates of most hymenopteran branches are relatively comparable, but we also noticed several notable accelerations of the gene gain and loss rates in different lineages. For example, the clade comprising *Nasonia*, *Trichomalopsis*, and *Pteromalus* in Pteromalidae exhibited an average gene gain and loss rate of 605.74 per My. Furthermore, a noticeable acceleration in gene gain and loss rates can be observed in the branches of leafcutter ants (Formicidae), as previously noted in studies^12,13^ (Supplementary Figs. 5–6 and Supplementary Table 4).

We identified 230 gene families that exhibited significant expansions or contractions during Hymenoptera evolution, and GO (Gene Ontology) enrichment analysis showed that they were enriched for the steroid metabolic process, the maltose metabolic process, and sensory perception of smell (FDR-adjusted *P* < 0.05; Supplementary Tables 5–6 and Supplementary Fig. 7). Among these, several rapidly evolved digestion-related gene families, such as Maltase, Esterase, and Trypsin, may be associated with adaptation to different diets in Hymenoptera (Supplementary Table 5). The rapid evolution of the Fatty acyl-CoA reductase family involved in the biosynthesis of insect pheromones may be driving the evolution of pheromone diversity^14–16^. The significant expansions and contractions of Odorant receptor families may be related to the rapid adaptations to detect the different odors from the living environments (such as pheromones and odors related to host plants or host insects) of diverse hymenopterans. We also found that the size of the cuticular protein family changed rapidly during the Hymenoptera evolution (Supplementary Fig. 8), which might be associated with the evolutionary diversity of the cuticular exoskeleton and morphology^17,18^. Detailed analysis further indicated that the size variation of the cuticular protein family mainly occurred in the CPR-RR-1, CRP-RR-2, and CPAP1 subfamilies (Supplementary Table 7 and Supplementary Fig. 8).

We also investigated the evolution of protein domain families based on the Pfam annotations. Similarly, we identified 105 Pfam domain families with significantly accelerated rates, including OS-D (Insect pheromone-binding family, A10/OS-D, PF03392.12), FA_desaturase (Fatty acid desaturase, PF00487.23), Chitin_bind_4 (Insect cuticle protein, PF00379.22), 7tm_2 (7 transmembrane receptor, PF00002.23), and Lipase (Lipase, PF00151.18), consistent with the significantly changed gene families (Supplementary Table 8). Furthermore, we reconstructed the rearrangements of protein domains in Hymenoptera evolution (see Methods). In total, 33,943 domain arrangement changes were observed, and 41.8% of them were formed by a fusion of two ancestral domains (Supplementary Table 9). We noticed the higher rates of domain rearrangements in several lineages, such as *G. flavifemur*, *Temnothorax longispinosus*, and *Diadromus collaris*, which may contribute to the evolutionary innovations of these species (Supplementary Table 9 and Supplementary Fig. 9).

We next explored the changes in the evolutionary rate of the protein-coding genes in Hymenoptera. We first estimated the rates of amino acid substitutions across the Hymenoptera phylogeny to reveal the overall pattern of protein sequence evolution (Fig. 1). Our analysis estimated an average amino acid substitution rate of 0.0025 substitutions per site per My with a standard deviation of 0.0055 (Supplementary Table 10). We found that the overall accelerated protein evolution events were dispersed across the multiple lineages of Hymenoptera, such as some clades in Anthophila (Apoidea) and Braconidae (Ichneumonoidea). These evolutionary rate shifts may be related to phenotypic transitions. For example, a comparative genomic study has suggested that the accelerated evolution of a number of proteins is linked to the independent miniaturization across parasitoid wasps^19^. Then, we examined the evolutionary rate of each orthologous protein-coding gene by measuring both the normalized amino acid substitution rates and the selective constraint (ratio of non-synonymous to synonymous substitutions, d_N_/d_S_). Functional analysis showed that genes associated with gene expression evolved most rapidly, followed by genes related to immunity, metabolism, and sensory perception (Fig. 2B). The rapid evolution of genes in these functional categories may play a role in phenotypic innovation and adaptive evolution in Hymenoptera. In contrast, genes with slower evolutionary rates were mainly enriched in some housekeeping processes, such as cellular component organization and cell development (Supplementary Fig. 10). In addition, we also conducted whole genome alignment analysis (Cactus alignment) to investigate the evolutionary constraints and acceleration at the whole genome level (see Methods). We chose the bumblebee *Bombus terrestris* (Apidae) as a reference because the availability of ATAC-Seq data is helpful to infer potential regulatory elements^20^. In the 27-way alignment, 59.5% (233.69/392.96 Mb) of the *B. terrestris* genome was aligned to at least one species, and 21.5% and 8.5% of the genome were identified as constrained and accelerated regions, respectively, by calculating the phyloP score (Supplementary Fig. 11A). We then identified the top 5% most accelerated and most conserved genes, measured by the average phyloP score of the coding sequence of each gene (Supplementary Table 11). GO enrichment analyses indicated that the most constrained genes were enriched in structure morphogenesis and development, while the most accelerated genes were involved in sensory perception and steroid metabolic processes, consistent with our previous findings in Fig. 2B (Supplementary Figs. 11B–C and Supplementary Tables 12– 13). We also identified 481,924 highly conserved elements using phastCons, and 27,602 (5.7%) of them were conserved across all species in our 27-way alignment (Supplementary Fig. 12A). Among them, 151 non-coding regions were overlapped with the accessible chromatin regions supported by ATAC-Seq in the reference species *B. terrestris*, suggesting their potential functions in gene regulation. We further found that the genes located near these conserved non-coding elements were functionally related to nervous system development and cell morphogenesis (FDR-adjusted *P* < 0.05; Supplementary Fig. 12B and Supplementary Table 14).

**Fig. 2.**
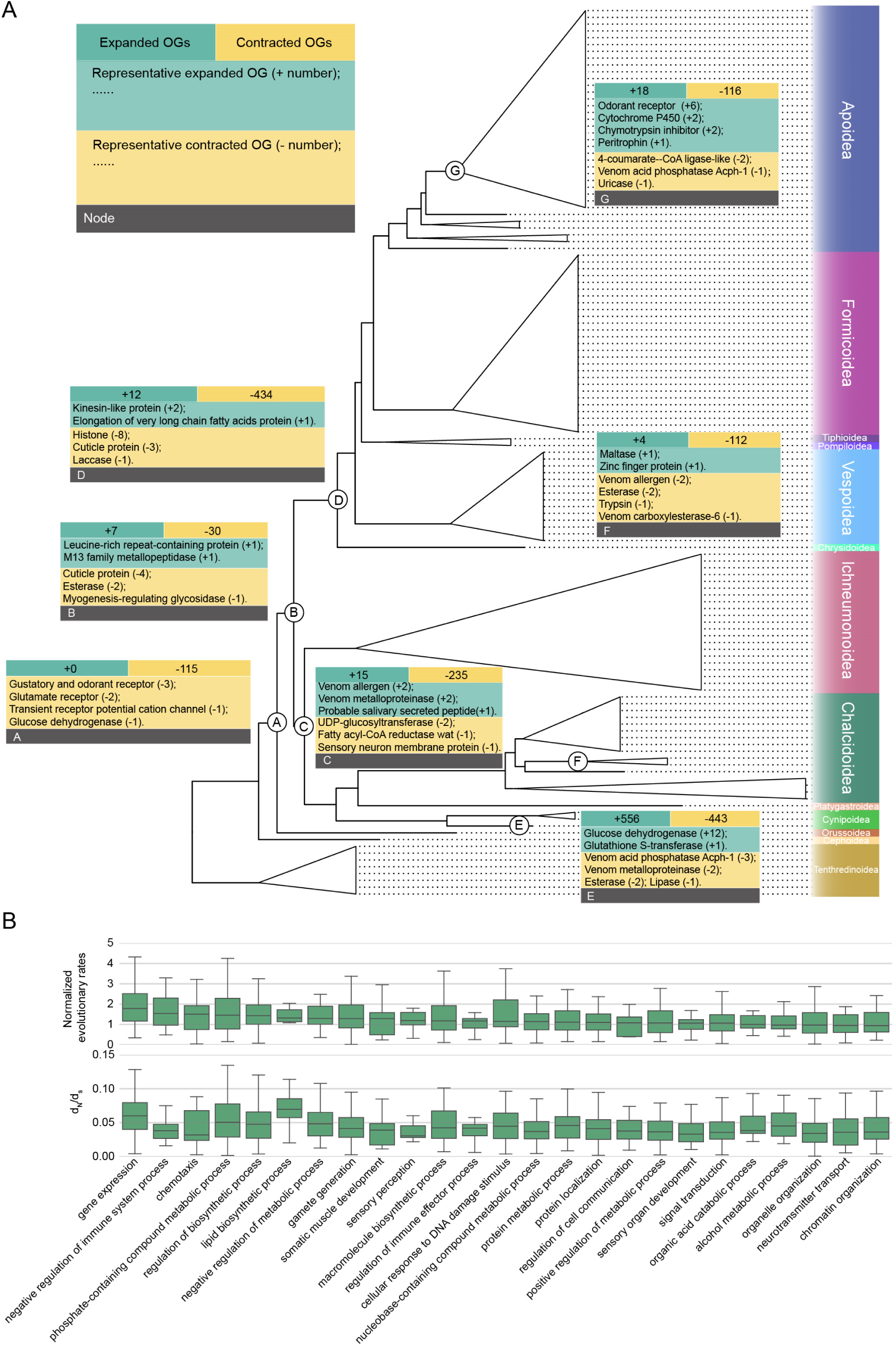
Gene family evolution and rapidly evolving genes in Hymenoptera. **A**) Gene family expansion and contraction at the nodes of key innovations in Hymenoptera evolution. In each box, the first row shows the number of gene families that expanded or contracted, respectively. Below are the corresponding representative gene families, and the size changes of each gene family are also indicated. **B**) Rapidly evolving OGs and functional categories in terms of evolutionary rate and selection constraint (d_N_/d_S_). Functional categories (GO Biological Process terms) are sorted by normalized evolutionary rates, and the top 25 GO terms are shown. The normalized evolutionary rate of each OG is measured as the mean of the amino acid substitution rates of each OG normalized by the genome-wide amino acid substitution rates. OGs were further grouped according to their functional similarity. d_N_/d_s_ of each OG refers to the ratio of the non-synonymous substitution rate to the synonymous substitution rate, measured by the one-ratio model in codeml.

### Genomic characteristics accompanying key innovations in Hymenoptera evolution

We next examined several key nodes in the evolutionary history of Hymenoptera, investigating the genomic changes that occurred at these nodes with the aim of correlating the association of genomic changes and phenotypic innovations. Here, we focused on four key evolutionary positions: the origin of parasitoidism (node A in Fig. 2A), the emergence of the wasp-waist (node B), the stem of Parasitoida that is prior to the radiation of the majority of parasitoid wasps (node C), and the emergence of the stinger (node D). The genomic features related to secondary phytophagy in several independent lineages will be discussed in a subsequent section. We did not investigate the genome evolution of eusocial hymenopterans due to the significant contributions of some recent works in this area^21–26^.

We first focused on the last common ancestor of the parasitoid wood wasp *Orussus abietinus* (Orussidae) and other wasp-waisted species (Apocrita), which represents the position of parasitoidism origin in the Hymenoptera tree. Our analysis identified 115 contracted and no expanded gene families at this node (Fig. 2A and Supplementary Table 15). Furthermore, we also found 137 missing gene families (that is, gene families present in over 70% of outgroup species and are lost in the test group) and only five novel gene families (also known as novel core genes, which are retained in over 70% of species in the test group) (Supplementary Tables 16–17). This pattern of gene family evolution suggests that the reductive evolution of gene families may have played an important role in the origin of parasitoidism. GO enrichment analysis showed that these contracted and missing families were significantly enriched for functions associated with detection of stimulus, cholesterol transport and metabolism, lipid metabolism, and xenobiotic metabolic processes (FDR-adjusted *P* < 0.05; Supplementary Tables 18–19).

The changes in these gene families in the ancestral parasitoid wasp could be associated with the shift in lifestyle and diet from a phytophagous sawfly to an insectivorous parasitoid. For instance, the genes related to detection of stimulus (e.g., Gustatory and odorant receptors, Glutamate receptors, and Transient receptor potential channels) may contribute to early parasitoid wasps locating potential hosts in novel environments (i.e., in dead wood for the most parasitoids in Orussidae^11^). The contracted gene families with cholesterol-transporting and metabolizing functions may be associated with alterations in the nutritional composition of the foods consumed after the acquisition of parasitoidism. In addition, we identified 96 genes that show evidence of positive selection in the parasitoid ancestor (aBSREL, FDR-adjusted *P* < 0.05; Supplementary Table 20). These genes were associated with many developmental processes, including muscle tissue development, eye morphogenesis, respiratory system development, and larval salivary gland morphogenesis (Supplementary Table 21). These genomic changes may be related to the dramatic changes in morphological characteristics that are considered adaptations to a parasitoid lifestyle, such as simplified larval morphology lacking eyes and legs in Orussidae^11,27^.

In the last common ancestor of wasp-waisted species (Apocrita), we also identified more contracted gene families (30) than expanded gene families (7) (Fig. 2A and Supplementary Table 22). The contraction of Cuticle protein genes that have roles in cuticle development at the Apocrita stem may contribute to the emergence of wasp-waist. We also discovered 119 positively selected genes with diverse functions such as endocytosis and regulation of enzyme activity (aBSREL, FDR-adjusted *P* < 0.05; Supplementary Table 23). At the stem of stingers, the Aculeata, 434 contracted gene families were found, but only 12 expanded gene families were found (Supplementary Table 24). The expanded genes included the Elongation of very long chain fatty acid protein genes that are involved in female pheromone biosynthesis^28,29^. The contracted genes with functions related to cuticle development (e.g., Cuticle protein genes and Laccase genes) were also identified. Moreover, our analysis revealed 175 positively selected genes at the stem of the Aculeata (Supplementary Table 25). These included the genes that regulate the highly conserved pathways for development, such as Wnt signaling pathway^30^ (e.g., *FZD2*, *LRP6*, and *MLLT3*) and Notch signaling pathway^31^ (e.g., *Ift172*, *NCSTN*, and *POFUT1*). Changes in these genes may have contributed to the unique morphological features of stinging hymenopterans.

The Parasitoida includes a large number of parasitoid wasp species, and the stem of the Parasitoida clade showed 235 contracted and 15 expanded gene families (Fig. 2A and Supplementary Table 26). We found the expansion of Venom allergen genes and Venom metalloproteinase genes, which may contribute to venom functions to enhance the probability of parasitic success. Additionally, we inferred that the expanded salivary secreted peptide family may also play a role in feeding on hosts and regulation of host immunity, based on current knowledge of the functions of the salivary proteins in parasitoid wasps^32–34^. This result suggests that the expansion of gene families involved in parasitoid-host interaction occurred prior to the radiation of parasitoid wasps. Taken together, our findings provide a number of candidate genes that may be linked to the evolution of key innovations in Hymenoptera.

### Divergent trajectories of genomic change between Parasitoida and Aculeata

The Parasitoida and Aculeata clades diverged from the ancestral wasp-waisted wasp at around 243 Mya, representing two major groups of hymenopterans with clear differences in lifestyle, morphology, and diversity^3^ (Fig. 1). The Parasitoida clade includes primarily parasitoid wasps that lack a stinger, and the Aculeata clade comprises stinging species with diverse life habits, including stinging parasitoids, social wasps, ants, and bees^3,8^. We next sought to explore and compare the genomic changes within these two clades, which may provide insights into the independent evolution of two major groups of hymenopterans.

Depending on the results of gene family evolution, we determined the distribution of the rapidly evolving events for each gene family on the branches in the Parasitoida and Aculeata clades. This thus allowed us to explore whether there are gene families with a clear distributional preference for rapid evolutionary events in the two clades. A total of 91 gene families were identified that exhibited rapid evolutionary events, with a significantly higher prevalence observed in the Parasitoida compared to Aculeata (FDR-adjusted *P* < 0.05, Chi-square test, odds ratio > 1; Fig. 3A, Supplementary Fig. 13, and Supplementary Table 27). These included a number of Histone genes (i.e., H2A, H2B, H3, and H4) that are involved in the structure of chromatin^35,36^, suggesting the potential distinct chromatin structures in parasitoid wasps. The enrichment of rapidly evolving Trypsin, Glucose dehydrogenase, and UDP-glucosyltransferase families in the Parasitoida may be associated with the evolutionary adaptations of digestion and detoxification metabolism, enabling the parasitoid wasps to adapt to diverse arthropod hosts. The rapid evolution of the Peptidoglycan recognition protein family in the Parasitoida may contribute to the antimicrobial defenses of parasitoid wasps^37,38^. Furthermore, we observed significant changes in the SET and MYND domain-containing protein family, an epigenetic regulator^39^, in the Parasitoida, which suggests that epigenetic regulation may play an important role in the evolution of parasitoid wasps. By contrast, only 12 gene families were identified as enriched rapidly evolving families in the Aculeata, including the THAP domain-containing protein, RCC1 and BTB domain-containing protein, Odorant receptor, and Gustatory receptor (FDR-adjusted *P* < 0.05, Chi-square test, odds ratio > 1; Supplementary Table 28 and Supplementary Fig. 14). In addition, we estimated and compared the rates of gene gain and loss for each gene family between the Parasitoida and Aculeata clades to determine the gene families with significantly different gene gain and loss rates within the two clades. This analysis may capture additional gene families that have undergone significant evolutionary changes between the Parasitoida and Aculeata, which may have been overlooked in previous analyses due to the limited number of evolutionary branches where such changes occurred. In this analysis, we identified 1604 gene families with significantly higher rates of gene gain and loss in the Parasitoida compared to Aculeata (FDR-adjusted *P* < 0.05, One-tailed Mann-Whitney *U* test; Fig. 3B and Supplementary Table 29). GO analysis of the gene families with high turnover rates in the Parasitoida showed enrichment for functional categories such as polysaccharide digestion, intracellular sterol transport, and prostaglandin secretion (FDR-adjusted *P* < 0.05; Fig. 3C and Supplementary Table 30). Additionally, we noticed some genes related to venom functions were included in this gene set, suggesting venom may play a key role in the evolution and diversity of parasitoid wasps in Parasitoida (Fig. 3B; discussed in detail in the next section). Similarly, we found a small number of gene families (26) with faster gene gain and loss rates in the Aculeata compared to Parasitoida (FDR-adjusted *P* < 0.05, One-tailed Mann-Whitney *U* test; Supplementary Table 31).

**Fig. 3.**
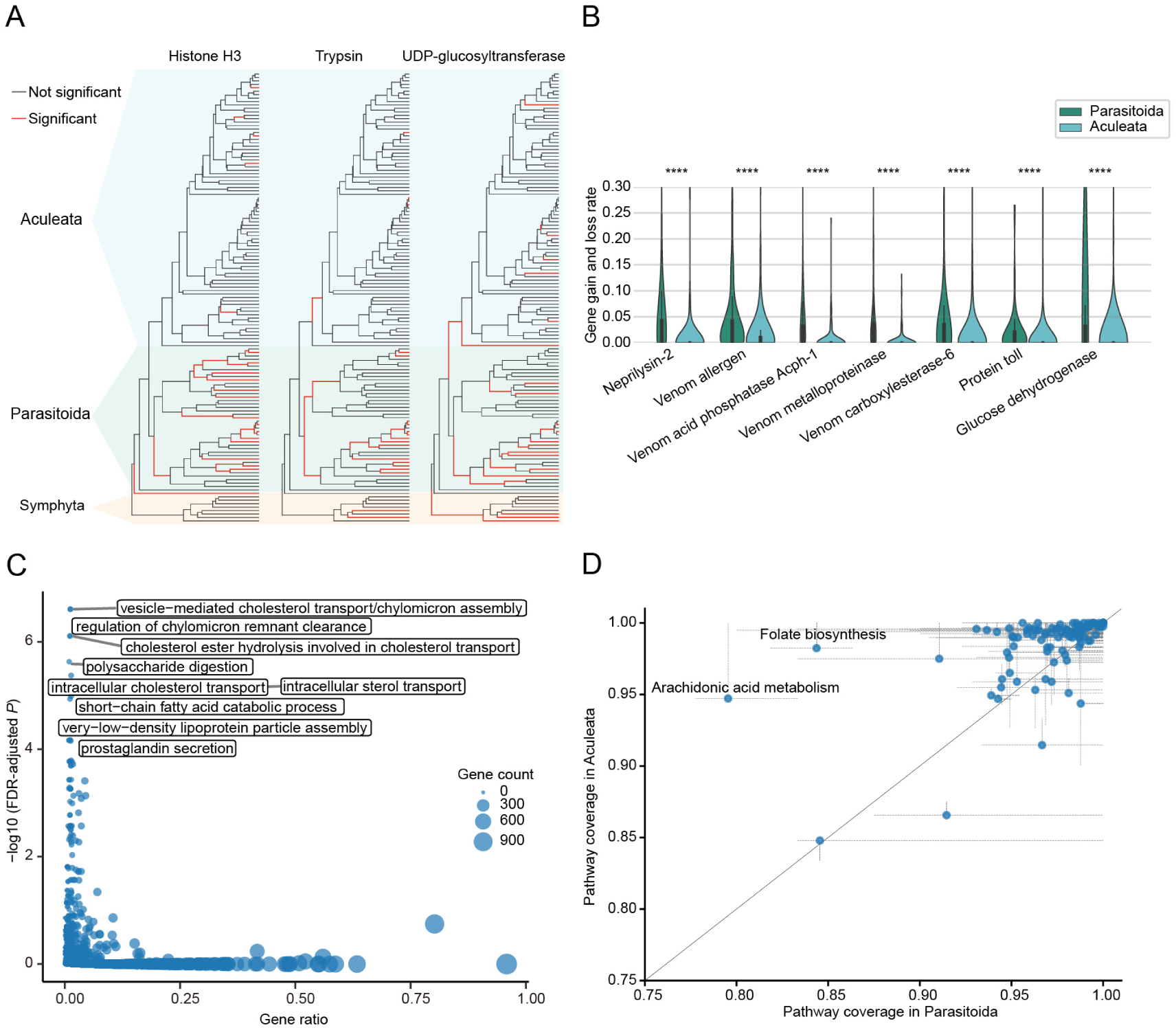
Different evolutionary genome dynamics between Parasitoida and Aculeata. **A**) The distribution of the rapidly evolving events for gene families on the branches of the Parasitoida and Aculeata clades. Three gene families, Histone H3, Trypsin, and UDP-glucosyltransferase, are shown here, which have a significantly higher occurrence rate of rapid evolutionary events in Parasitoida than in Aculeata (FDR-adjusted *P* < 0.05, Chi-square test, odds ratio >1). **B**) Gene families that have a higher rate of gene gain and loss in Parasitoida than in Aculeata. The gene gain and loss rate is measured as the sum of gene gains and losses per million years. Asterisks indicate significant shifts between Parasitoida and Aculeata (One-tailed Mann-Whitney *U* test; *≤05; **≤0.01; ***≤0.001, ****≤0.0001). **C**) GO enrichment analysis of the gene families with a faster rate of gene gain and loss in Parasitoida than in Aculeata. The gray box shows the GO terms for the 10 biological processes with the lowest adjusted *P* value (FDR-adjusted *P* < 0.05, Benjamini-Hochberg multi-test correction). The size of the blue circles represents the number of genes in the corresponding GO. **D**) KEGG pathway coverage comparison between Parasitoida and Aculeata. The expanded lines from the point represent the first and third quartiles of pathway coverage in Parasitoida and Aculeata, respectively.

We next compared the selective constraint (d_N_/d_S_) of single-copy OGs between the Parasitoida and Aculeata. The result showed none of the OGs had significantly different d_N_/d_S_ between the two clades (Two-tailed Mann-Whitney *U* test; Supplementary Table 32). We hypothesized that this result may be attributed to the presence of significant variability in the d_N_/d_S_ ratio of the branches within the two clades. In order to investigate the degree of variation in d_N_/d_S_ between the Parasitoida and Aculeata, we calculated the coefficient of variation (CoV) of the d_N_/d_S_ in the two clades (Supplementary Table 33). Our findings revealed that the d_N_/d_S_ variation of the Parasitoida was significantly larger than that of the Aculeata (*P* = 7.92e-84, One-tailed Mann-Whitney *U* test; Supplementary Fig. 15). And 73.8% of the analyzed OG exhibited a greater CoV of the Parasitoida than that observed in the Aculeata (Supplementary Table 33). This result suggests that the Parasitoida may be subject to more variable selection pressures compared to the Aculeata. This is likely to be related to the diverse lifestyles and hosts of the parasitoid wasps in Parasitoida. We also identified 13 OGs with obviously higher CoV of the d_N_/d_S_ in the Parasitoida (i.e., the OGs with the top 10% CoV in Parasitoida and the bottom 10% CoV in Aculeata), including exocytosis-related STXBP5, calcium-binding protein Calmodulin, and cytoskeleton organization-related DIAPH, suggesting that they may have evolved under different selective pressures and may play key roles in Parasitoida evolution (Supplementary Table 34).

We further explored the variation in metabolic pathways between the Parasitoida and Aculeata, which may reflect differences in the life history of these two groups. To compare the pathway repertoire variation between the two clades, we first performed EC (Enzyme Commission number) annotations for all 131 hymenopterans (Supplementary Table 35) and constructed ancestral KEGG (Kyoto Encyclopedia of Genes and Genomes) pathways based on eight phytophagous sawflies (Tenthredinoidea) located on the early branches of the Hymenoptera tree (see Methods). Then, we mapped the present/absent information of each EC of the species from the Parasitoida and Aculeata to the ancestral pathways. This allows us to study the pathway-level changes of Parasitoida and Aculeata after their divergence. We examined the pathway coverage (PC, i.e., the fraction of ECs in an ancestral pathway that were annotated in a species) of each pathway in each species (Supplementary Table 36). Overall, our results showed significantly lower PC for Parasitoida compared to Aculeata (*P* = 2.68e-46, One-tailed Mann-Whitney *U* test; Supplementary Fig. 16), suggesting that the specialized parasitoidism may have simplified the pathways of parasitoid wasps, and similar phenomena have also been demonstrated in other parasitic species, such as parasitic worms^40^, dicyemids^41^, protists^42^, and plants^43^. Our detailed analysis further revealed that 37 pathways from 14 KEGG superpathways exhibited significantly lower PC in Parasitoida when compared to Aculeata, which are involved in lipid metabolism, metabolism of cofactors and vitamins, carbohydrate metabolism, etc. (FDR-adjusted *P* < 0.05, One-tailed Mann-Whitney *U* test; Fig. 3D, Supplementary Fig. 17, and Supplementary Table 37). Conversely, five pathways showed lower PC for Aculeata, and these metabolic processes included purine metabolism, pentose phosphate pathway, and glyoxylate and dicarboxylate metabolism (FDR-adjusted *P* < 0.05, One-tailed Mann-Whitney *U* test; Fig. 3D, Supplementary Fig. 17, and Supplementary Table 38). Among these PC variations, we inferred the lower PC of arachidonic acid metabolism (ko00590) and folate biosynthesis (ko00790) in Parasitoida caused by the pervasive absence of carbonyl reductase (K00079, EC. 1.1.1.184/189/197). In addition, we found that the loss of the same enzyme may have different effects on these two pathways. Specifically, the absence of carbonyl reductase in the arachidonic acid metabolic pathway has disrupted the pathway from prostaglandin E2 to prostaglandin F2alpha. The widespread absence of this pathway in parasitoid wasps suggests that they may have other ways of acquiring prostaglandin F2alpha, probably from their diet. However, the absence of carbonyl reductase in the folate biosynthetic pathway did not result in pathway disruption due to the redundancy in this pathway. This may represent a streamlined evolution of the pathway in parasitoid wasps. We next compared the CoV of the PC between the two clades and found six superpathways exhibited significantly higher variation in Parasitoida, also covering the pathways we discussed above (FDR-adjusted *P* < 0.05, One-tailed Mann-Whitney *U* test; Supplementary Fig. 18). This finding suggests that these pathways are more variable in parasitoid wasp species, and it is likely that these variations occurred several times independently in Parasitoida evolution. This may suggest that the rapid changes in these pathways are related to the adaptation of parasitoid wasps to diverse parasitoid lifestyles, such as adaptation to different hosts and different parasitic strategies.

### Massively duplicated genes and gene losses in Parasitoida

The preceding section of this study has demonstrated that parasitoid wasps exhibit high rates of gene gain and loss, which likely results in significant changes in the number of members of certain gene families. Here we first explored the gene families with significantly larger gene numbers in Parasitoida than those in other taxa, which were likely to be caused by the massive gene duplications that occurred in Parasitoida. In total, we identified 95 gene families with a significantly larger number of genes in Parasitoida species compared to other hymenopterans (FDR-adjusted *P* < 0.05, One-tailed Mann-Whitney *U* test; Supplementary Table 39 and Supplementary Fig. 19). Functional annotation of these families was diverse, but we observed they were frequently related to chemoreception (such as Odorant receptor, Gustatory receptor, and General odorant-binding protein), detoxification (such as Multidrug resistance-associated protein and UDP-glycosyltransferase), and protein degradation (such as Proteases and Protease inhibitors). Among these gene families, we also found 41 families with no annotation information, designated as Putative uncharacterized proteins. In contrast, we only found nine enlarged gene families for Aculeata, with three families belonging to the Odorant receptor (Supplementary Fig. 20 and Supplementary Table 40). The findings suggest that large-scale gene duplication was more prevalent in Parasitoida, and the identified gene families that exhibited massive expansions may have played roles in the adaptive evolution of parasitoid wasps and promoted the diversity of these species.

Among the gene families identified with large-scale gene duplications in Parasitoida, we found several families whose members are often known as venom components, the important weapons of parasitoid wasps to overcome host immune systems or manipulate host development^44,45^. For instance, our analysis indicated that the Venom metalloproteinase family has undergone an extensive expansion, specifically in the Parasitoida clades, which may have contributed to the recruitment of the family member as a venom component in parasitoid wasps through neofunctionalization (Fig. 4A and Supplementary Fig. 21). Indeed, Venom metalloproteinase has been reported as venom protein in at least 19 parasitoid wasps from 5 families of 3 superfamilies in Parasitoida^46–62^. And some studies have demonstrated that these venom proteins play distinct roles (such as regulating host development^57^ and inhibiting host immunity^63^) in different parasitoid-host systems, suggesting the duplications of this family in parasitoid wasps may also be related to the diversity of venom functions. Additionally, the duplication of Venom metalloproteinase genes may also help diversify nonvenomous functions. In *Anastatus fulloi* (Chalcidoidea, Eupelmidae), our gene expression profiling showed diverse expression patterns of Venom metalloproteinase genes, suggesting that they have functions other than venom functions (Fig. 4B). For example, nine of them exhibited specifically higher expression in the larval stage, suggesting their possible roles in digestion or parasitoid-host interaction as salivary proteins (Fig. 4B and Supplementary Table 41). Similarly, the expanded Venom allergen family in Parasitoida may also be related to the venom evolution of parasitoid wasps (Fig. 4C and Supplementary Fig. 22). And we found 15 parasitoid species have used the members of this family as their venom^47,53,54,58–61,64–69^. Furthermore, the gene family also exhibited diverse gene expression patterns, which suggests that the gene may have multiple functions (Fig. 4D and Supplementary Table 42). Other enlarged gene families in Parasitoida that have evidence related to venom components include Neprilysin^70,71^, Serpin^53,72,73^ and Trypsin^62^. Overall, our findings indicate that the extensive duplication of certain genes in parasitoid wasps has played a role in the evolution of venom diversity. This may enhance the adaptability of parasitoid wasps to different hosts, thereby contributing to the species diversity in Parasitoida.

**Fig. 4.**
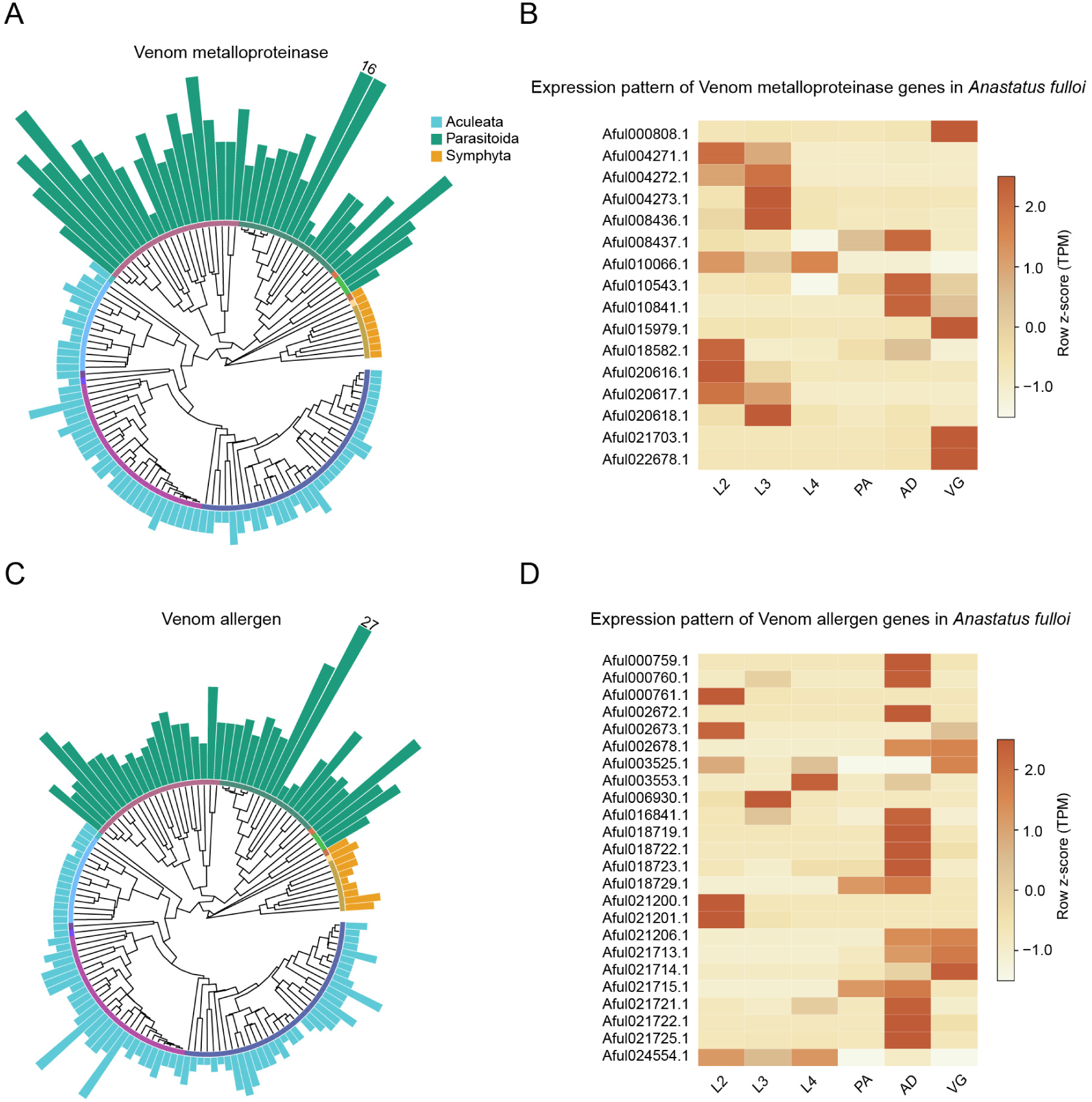
Expansions of Venom allergen and Venom metalloprotease gene families in Parasitoida. **A**) The size distribution of the Venom metalloprotease gene families in Hymenoptera. The number on the bar indicates the maximum gene copy count across the species for that family. **B**) Expression pattern of the Venom metalloprotease genes in *Anastatus fulloi* at five developmental stages and the venom gland. Expression is measured using z-score values of transcripts per million (TPM) by row. L2, second instar larva; L3, third instar larva; L4, fourth instar larva; YP, pupal stage; AD, female adult; VG, venom gland. Only genes with a TPM greater than 0 in at least one period are shown. **C**) The size distribution of the Venom allergen gene families in Hymenoptera. **D**) Expression patterns of the Venom allergen genes in *A. fulloi* at five developmental stages and the venom gland.

We also identified 199 gene families that were completely missing in a higher proportion of Parasitoida than in other taxa (i.e., the gene family is lost in more than 20% of parasitoids but in no more than 2% of other taxa) (Supplementary Table 43). Several losses in these gene families may be associated with the parasitoid lifestyle of species in Parasitoida. For example, we hypothesized that the loss of some gene families/subfamilies associated with detoxification metabolism (such as UDP-glycosyltransferase, Cytochrome P450, and Nephrin) may be related to the reduction of toxic substances contained in the hosts of some parasitoids. Eleven gene families with GO terms related to embryo development exhibited a higher loss proportion in parasitoid wasps (Supplementary Table 43). This suggests that the early developmental processes of parasitoid wasps may have been altered compared to other hymenopterans, possibly in response to adaptations to parasitic life. In addition, our analysis also recovered our previous finding that Vitellogenin and its receptor (Vitellogenin receptor) have been repeatedly lost in many parasitoid wasps^74^. The loss of these genes may result in a reduction or absence of yolk proteins in parasitoid wasp eggs, which is consistent with some phenotypic observations^75–77^. It is proposed that this may be related to the fact that parasitoid wasp eggs are able to obtain nutrients from the host.

### Genomic features of independent secondary phytophagy

During the evolution of Hymenoptera, a shift from phytophagy to parasitoidism occurred at the ancestral node of Orussidae and Apocrita, after which some hymenopterans from different superfamilies re-evolved into phytophagy (i.e., secondary phytophagy). And a recent evolutionary study has proposed that secondary phytophagy in Hymenoptera plays a key role in their species diversity^4^. However, the genomic basis for the hymenopterans that evolved back to phytophagy is still unknown. Moreover, it is worth exploring whether there are any similarities in the genomic features underlying these independent transitions (i.e., convergent or parallel evolution). In the present study, our Hymenoptera phylogeny captured three independent transitions to secondary phytophagy, locating at the terminal branch leading to the gall wasp *Belonocnema kinseyi* (Cynipoidea, Cynipidae) (node E in Fig. 2A), the ancestral branch of two fig wasps *Ceratosolen solmsi* and *Eupristina verticillata* (Chalcidoidea, Agaonidae) (node F), and the ancestral branch of pollen-feeding bees (Apoidea, Anthophila) (node G; Fig. 1). First, we examined the genomic changes, including gene family expansions/contractions and genes with positive selection on these target branches. In general, the genomic changes of these three branches showed a high degree of branch specificity, both in terms of the number of changes and the composition of genes or gene families (Fig. 5A and Supplementary Tables 44–46). In addition, our analysis did not reveal any instances of gene family expansion, contraction, or positive selection of genes that were common to all three branches (Fig. 5A). This result suggests that the genomic changes behind the three independent transitions to phytophagy in Hymenoptera may be distinct. This may be attributed to differences in the types of plant-derived foods consumed (e.g., plant galls for the gall wasp and pollen for bees). Nevertheless, we did identify some genomic changes that were shared by two branches or specific to one branch, which may be associated with secondary phytophagy (Fig. 2A). For example, we observed that the Maltase family has expanded on the branches related to the secondary phytophagy of both the gall wasp and fig wasps, which probably play a role in enhancing the carbohydrate metabolism of these species (Supplementary Table 44). This is also supported by the expansion of Glucose dehydrogenase family in the gall wasp *B. kinseyi*. We also noticed the family contractions (e.g., Venom metalloproteinase and Venom allergen) related to venom functions in the gall wasp and fig wasps (Supplementary Table 45). This may be indicative of the degradation of the venom components originally used to attack insect hosts following their transformation to phytophagy. In bees, a large expansion of Odorant receptors was observed, probably to improve the ability to recognize and distinguish volatile substances in plants so as to effectively recognize and select food. Additionally, we found the Peritrophin family expanded, which is regarded as the most important physical immune barrier of the gut in arthropods^78^, possibly as an adaptation from sarcophagy to pollen feeding. In contrast, the shrinkage of the Uricase gene could be an evolutionary response to the lower metabolic capacity for purine degradation^79^. In *B. kinseyi*, the expansion of Glutathione S-transferase could also be required for the detoxification of phytochemicals^80,81^ (Fig. 2A). Meanwhile, positive selection analysis resulted in 97, 73, and 148 positively selected genes in fig wasps, bees, and the gall wasp, respectively, of which 272 genes were exclusive (Fig. 5A and Supplementary Table 46). We further discovered that 67.28% of OGs to which exclusively positively selected genes belong were clustered into 19 functional clusters composed of OGs with genes evolving under positive selection exclusively in three transition branches, probably indicating the functional convergence signal (Fig. 5B and Supplementary Table 47). However, no more than 30% of expanded and contracted OGs could be clustered (Supplementary Figs. 23–24 and Supplementary Tables 48–49). Overall, the genomic signatures of dietary changes can be summarized into three functional categories: chemoreceptor, digestion, and detoxification.

**Fig. 5.**
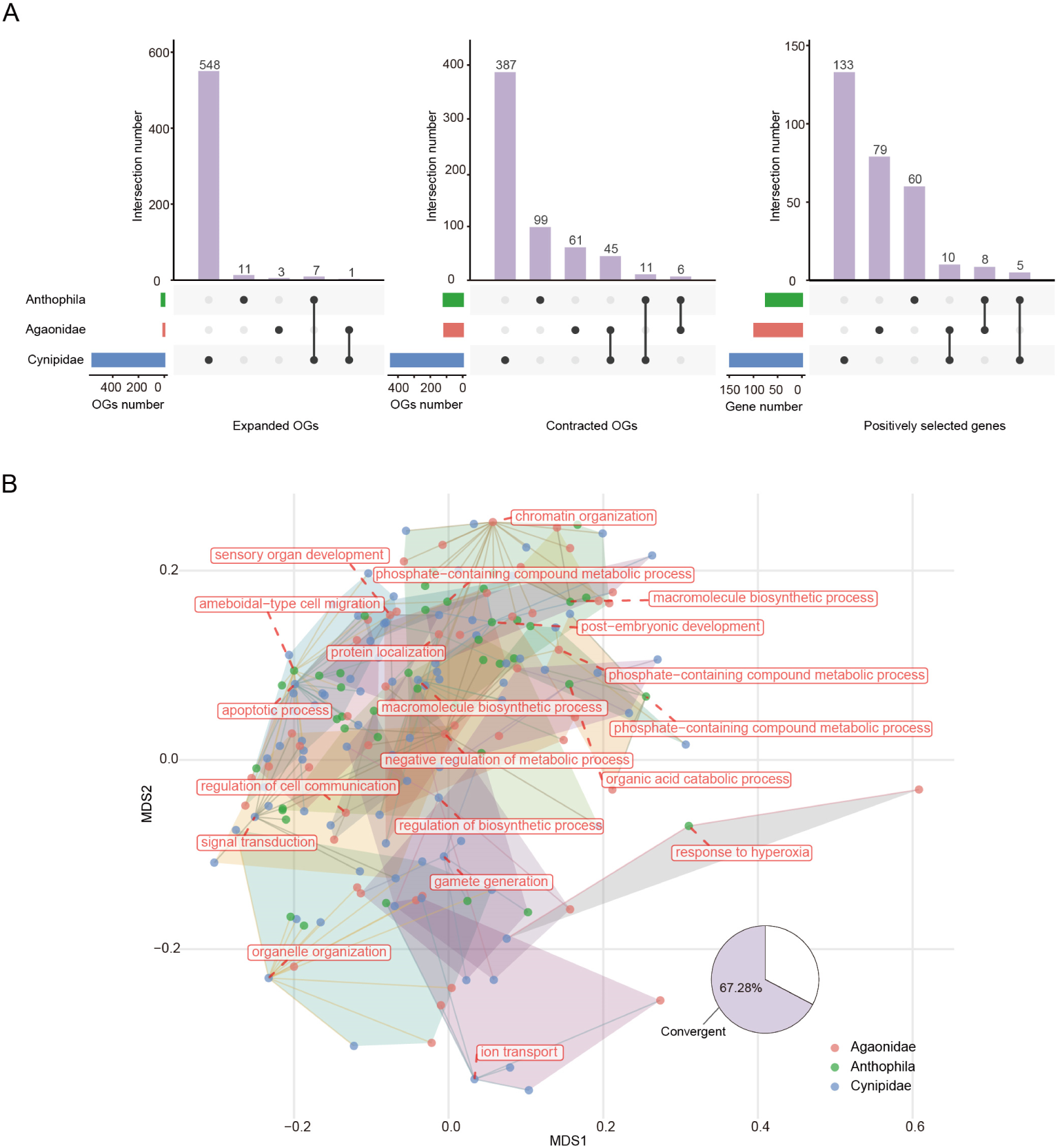
Genomic features of three independent origins of secondary phytophagy. **A**) UpSet charts showing the number of shared and unique expanded, contracted, and positively selected gene families among three clades of independent transitions to secondary phytophagy: Cynipidae, Agaonidae, and Anthophila, with intersection size representing the number of shared OGs in each category. No expanded or contracted OGs, or positively selected genes, were shared by all three groups. **B**) Functional convergence of exclusively positively selected genes in the three branches of secondary phytophagy origins. Each dot represents an OG to which the exclusively positively selected gene belongs. The centroid OG of each polygon is labeled with potential functions. Of the OGs with exclusively positively selected genes, 67.28% were clustered into 19 functional clusters, indicating functional convergence signals.

Plants exploit the glucosinolate-myrosinase system to produce toxic hydrolysis products to defend themselves from herbivores^82,83^. In *Athalia rosae* and some aphids, glucosinolate can be rapidly sequestered into the hemolymph, where it is hydrolyzed by myrosinases^11,84–86^. Thus, a high copy number of the myrosinase gene may characterize phytophagous species^11^. However, in our study, the copy number of myrosinase was indeed high in phytophagous sawflies, but there was no expansion in secondary phytophagous species (Supplementary Fig. 25). Therefore, we speculate that secondary phytophagous species may employ strategies of detoxification metabolism that are different from those of phytophagous species.

### Conclusion

In this study, our evolutionary and comparative genomic analyses revealed many genomic changes (such as gene family evolution, protein domain evolution, evolutionary rate, and selection) in the evolutionary history of Hymenoptera. We correlated some genomic changes with specific innovations, including the evolution of parasitoidism, wasp-waist, sting, and secondary phytophagy. In comparing the two major hymenopteran clades, Parasitoida and Aculeata, we uncovered distinct genomic features indicative of their independent evolutionary trajectories, including variations in pathway coverage and gene family dynamics, which may reflect adaptation of their divergent lifestyles (highly parasitoidism vs. complex free-living habits, which are not limited to parasitoidism). Furthermore, we investigated the genomic basis for the hymenopterans that evolved back to phytophagy in three independent lineages. Overall, the results of our study provide insights into the evolution and diversification of Hymenoptera.

## Methods

### Data preparation

We collected publicly available genomic and annotation data for 131 Hymenoptera species from 13 superfamilies and 29 families. To avoid redundancy, we selected a representative protein isoform (i.e., the isoform with the longest amino acid sequence) for each gene in each species. Assembly quality of each genome was assessed with QUAST 5.2.0^87^, and completeness was assessed with BUSCO 5.5.0^88^ (-m prot -l insecta_odb10). Finally, the average BUSCO score reached 97.26%. For more detailed information about species and public data repositories, see Supplementary Table 1. To reduce the impact of potential gene annotation errors, we corrected potential chimeric genes and broken genes. We first used Broccoli v1.2^89^ to identify OGs from all protein sequences of 131 species. Genes assigned to two or more OGs by Broccoli v1.2 were considered chimeric genes. Those that belonged to the same OG, were sequenced on the same strand on the same chromosome, showing low sequence similarity (less than 30% calculated using MAFFT v7.310^90^), and matching different portions of the longest homolog protein in *Nasonia vitripennis* were considered broken genes. Detailed correction procedures for chimeric and broken genes are referenced in a previous study^10^.

### Phylogenetic analysis

We identified 33,405 OGs from all protein sequences of 131 species with OrthoFinder 2.5.4^91^. The protein sequences of 1002 single-copy genes were aligned with MAFFT v7.310^90^ and concatenated into supergenes. Then, these sequences were used to construct a maximum-likelihood phylogenetic tree using IQ-TREE 2.0.3^92^ with 1000 ultrafast bootstraps. The best-fitting model estimated by ModelFinder^93^ was JTT+F+R10. Species divergence times were estimated with r8s1.81^94^ using 13 time points based on previous studies^3^. The time points were as follows: Hymenoptera: 281 Mya, Tenthredinoidea: 106–176 Mya, Orussoidea + Apocrita: 211–289 Mya, Apocrita: 203–276 Mya, Ichneumonoidea: 151–218 Mya, Chalcidoidea: 105–159 Mya, Cynipoidea: 69–125 Mya, Aculeata: 160–224 Mya, Apoidea: 128–182 Mya, Vespoidea: 114–180 Mya, Formicidae: 65–127 Mya, Apidae: 68–99 Mya, and Braconidae: 116–177 Mya.

### Gene family evolution

We used CAFE5^95^ to determine gene family expansion and contraction for each branch, using the result of OrthoFinder and a species-level phylogeny with divergence times as inputs. Gene families with a *P* value less than 0.05 were considered to have significant expansions and contractions. In order to estimate the differences in the distribution of rapid evolutionary events between Parasitoida and Aculeata, we used Chi-square tests, and if FDR-adjusted *P* < 0.05 and odds ratio > 1, the gene family was considered to have a significantly different occurrence rate of rapid evolutionary events between Parasitoida and Aculeata. The gene gain and loss rate for each clade was calculated as the sum of gene gains and losses for all OGs divided by the divergence time. For each OG, the gene gain and loss rates of different branches were calculated as the count change of OG between it and its parent clade divided by the corresponding divergence time.

### Cuticular protein family analysis

BLASTP analyses were performed against protein sequences of all species using cuticular protein sequences from hymenopteran species (i.e., *Apis mellifera*, *Microplitis mediator*, *P. puparum*, and *N. vitripennis*) as probes (-evalue 1e-5). Meanwhile, the genomes of all species were searched by TBLASTN with the above probes. The domains of proteins aligned with BLASTP and TBLASTN were predicted using hidden Markov model searches with HMMER v3.4 (https://github.com/EddyRivasLab/hmmer) against Pfam database (http://pfam.xfam.org/). For the cuticular proteins analogous to the peritrophins (CPAP) family, the CBM_14 domain (PF01607) was required, of which the CPAP1 subfamily had one and CPAP3 had three. For the cuticular proteins with Rebers and Riddiford (R&R) consensus (CPRs), the Chitin_bind_4 domain (PF00379) was required, and they were divided into RR-1 and RR-2 subfamilies according to the best hit. The CPF family was identified based on the Cuticle_3 domain (PF11018) and the Tweedle (TWDL) family based on the DUF243 domain (PF03103).

### Selection pressure analysis

We selected 27 representative species of the major groups of Hymenoptera and identified 2060 one-to-one orthologue protein-coding genes using OrthoFinder 2.5.4^91^; for detailed species, see Supplementary Table 50. HyPhy^96^ package was used to detect gene selection signals at key points in Hymenoptera evolution. aBSREL^97^ was used for detecting positively selected genes (FDR-adjusted *P* < 0.05, Bonferroni-Holm correction). To explore the selection pressure of genes in each clade, we calculated d_N_/d_S_ using PAML 4.9^98^ in the free-ratio model, assuming that the d_N_/d_S_ values of all branches in the phylogenetic tree are not equal. Mann-Whitney *U* tests were performed to identify OGs with a significant difference in d_N_/d_S_ between Parasitoida and Aculeata. To further test d_N_/d_S_ variation in Parasitoida and Aculeata, we used the CoV, calculated by dividing the standard deviation by the mean of d_N_/d_S_ in each OG in each group, and compared them using a Mann-Whitney *U* test.

### Protein domain analysis

We utilized HMMER 3.3.2 to annotate the protein domains in all species and calculated the number of domain families classified based on PfamA accession. CAFE5^95^ was used to identify significantly accelerated domain families in the same way that gene families were. The domain rearrangements were reconstructed using DomRates^99^ with PfamScan (https://github.com/aziele/pfam_scan) results as inputs.

### GO annotation and enrichment analysis

We annotated proteins from all species using eggNOG-mapper v2^100^ against the eggNOG5 database. Based on the GO annotation results, we performed GO enrichment analysis of the candidate gene set using GOATOOLS^101^ with the go-basic.obo database (http://geneontology.org/). The result of the *P* values for each GO term was corrected for an FDR < 0.05 (Benjamini-Hochberg multi-test correction).

### Molecular evolution analysis on functional categories

The package constellatoR (https://github.com/MetazoaPhylogenomicsLab/constellatoR) was used to assign GO annotations to each OG and cluster GOs with similar functions using the results of eggNOG-mapper v2 and OrthoFinder as inputs, thus classifying OGs into different functional categories. For each OG in each category, the normalized evolutionary rate was estimated by averaging the ratio of amino acid substitution rates of interspecific homologous proteins to inter-species amino acid substitution rates. PAML 4.9^98^ was used to calculate the d_N_/d_S_ ratio for each OG using its respective phylogenetic tree.

### Whole genome alignment and constraint analysis

We used a 27-way whole genome alignment generated by the reference-free aligner Cactus^102^ and PHAST v1.6^103^ programs (https://github.com/CshlSiepelLab/phast). We utilized msa_view to retrieve 4-fold degenerate sites based on *B. terrestris* coding gene annotation, and we fitted a phylogenetic model with phyloFit from PHAST v1.6. To quantify the constraint of individual nucleotides, we utilized phyloP (part of the PHAST v1.6 package) to compute the per-base constraint based on the resulting model. Columns with a positive score were considered constrained bases, and those with a negative score were accelerated-evolving bases. Similarly, we used phastCons^104^ (part of the PHAST v1.6 package) to find conserved elements. We excluded conserved elements less than 20 bp in length. To further investigate the chromatin accessibility of conserved no-coding elements, we downloaded raw ATAC-Seq data for the egg, larva, pupa, and adult stages of *B. terrestris*^20^ from NCBI (accessions in Supplementary Table 51). Trimmomatic 0.33^105^ was used to filter raw reads, removing low-quality reads and adaptor sequences. Clean reads were mapped to the *B. terrestris* reference genome with Bowtie2^106^. Then, we used Picard 3.1.1 (https://github.com/broadinstitute/picard) to eliminate PCR duplicates and SAMtools 1.6^107^ to delete mitochondrial reads. MACS2^108^ was utilized to identify peaks, and the irreproducible discovery rate (IDR) analysis was performed to assess the reproducibility by comparing the consistency of two biological replicates at the same developmental stage. Peaks with IDR <= 0.05 were reproducible across two replicates and were kept for joint analysis with identified conserved no-coding elements.

### Metabolic pathway evolution

We used eggNOG-mapper v2^100^ to perform EC annotations for all 131 hymenopterans, and then ancestral pathways were constructed according to the intersections of annotated ECs in eight sawflies (Tenthredinoidea) and the enzymes of each pathway in KEGG. Pathway coverage was measured as the number of ECs in the pathway annotated divided by the total number of ECs in the constructed ancestral pathway. We considered pathways for which KEGG had a reference pathway for *A. rosae*, and the reconstructed pathway contained at least five ECs. Coverage of each pathway between the Parasitoida and Aculeata species was compared by a Mann-Whitney *U* test, and the FDR-adjusted *P* of less than 0.05 was considered significant higher or lower. The CoVs of pathway coverage in Parasitoida and Aculeata were calculated as the standard deviation divided by the mean of pathway coverage in these two clades, respectively.

### Functional convergence analysis

To investigate the level of functional convergence of interested OGs in three secondary phytophagy transitional ancestral branches, the R package constellatoR was used to calculate the similarity based on the semantic similarity matrix between OGs and cluster them with an exemplar OG. We then used the plotConstellation function to visualize the cluster shared by the interested OGs of the three transition nodes.

### Transcriptome analysis

Raw transcriptome sequencing data for the second instar larva, third instar larva, fourth instar larva, female pupa, female adult, and venom gland of *A. fulloi* were downloaded from NCBI (accessions in Supplementary Table 52). Fastp 0.22.0^109^ was used for filtration and quality control, and Salmon 0.14.1^110^ was used to calculate the expression level of transcripts (measured as transcripts per million, TPM).

## Acknowledgments

This work was supported by the Program of the National Natural Science Foundation of China (NSFC) (Grant No. 32202376 to XHY), the Key Program of NSFC (Grant No. 32330085 to GYY), the Young Elite Scientists Sponsorship Program by China Association for Science and Technology (Grant No. 2022QNRC001 to XHY), the Program of NSFC (Grant No. 32302408 to YY, No. 32302337 to XXZ, No. 32302444 to JLW), and the Scientific Research Startup Fund Project of Zhejiang A&F University (Grant No. 2023LFR119 to XHY). We are grateful to all colleagues who have contributed to the sequencing of Hymenoptera genomes.

## Author contributions

Project conception: X.H.Y. Manuscript preparation: C.H., Y.Y., X.X.Z., X.H.Y., and G.Y.Y. Data collection and preparation: C.H., Y.Y., L.J.P., Y.Y.L., S.X., Y.M., X.K.G., and H.X. Phylogenetic analysis: C.H. and Y.Y. Comparative genomic analysis: C.H., X.H.Y., Y.Y., X.X.Z., J.J.L., and Y.T.C. Data interpretation: X.H.Y., C.H., Y.Y., S.J.X., J.L.W., Z.C.Y., Z.W.T., Q.F., and G.Y.Y. All authors read and approved the manuscript.

## Competing interests

The authors declare no competing interests.

## Notes

### Competing Interest Statement

The authors have declared no competing interest.

### Summary of Updates

We revised Figures 1-2, and we updated the Methods section to make it clearer.

## References

1. Grimaldi, D. & Engel, M. S. Evolution of the Insects. (Cambridge University Press, 2005).

2. Huber, J. T. Biodiversity of Hymenoptera. in Insect Biodiversity: Science and Society (eds. Foottit, R. G. & Adler, P. H.) 303–323 (Wiley, 2009).

3. Peters, R. S. et al. Evolutionary history of the Hymenoptera. Curr. Biol. 27, 1013– 1018 (2017).

4. Blaimer, B. B. et al. Key innovations and the diversification of Hymenoptera. Nat. Commun. 14, 1212 (2023).

5. Sharkey, M. J. Phylogeny and classification of Hymenoptera. Zootaxa 1668, 521– 548 (2007).

6. Branstetter, M. G. et al. Phylogenomic insights into the evolution of stinging wasps and the origins of ants and bees. Curr. Biol. 27, 1019–1025 (2017).

7. Tang, P. et al. Mitochondrial phylogenomics of the Hymenoptera. Mol. Phylogenet. Evol. 131, 8–18 (2019).

8. Polaszek, A. & Vilhemsen, L. Biodiversity of hymenopteran parasitoids. Curr. Opin. Insect Sci. 56, 101026 (2023).

9. Vilhelmsen, L., Mikó, I. & Krogmann, L. Beyond the wasp-waist: structural diversity and phylogenetic significance of the mesosoma in apocritan wasps (Insecta: Hymenoptera). Zool. J. Linn. Soc. 159, 22–194 (2010).

10. Mantica, F. et al. Evolution of tissue-specific expression of ancestral genes across vertebrates and insects. *Nat*. Ecol. Evol. 8, 1140–1153 (2024).

11. Oeyen, J. P. et al. Sawfly genomes reveal evolutionary acquisitions that fostered the mega-radiation of parasitoid and eusocial Hymenoptera. Genome Biol. Evol. 12, 1099– 1188 (2020).

12. Wissler, L., Gadau, J., Simola, D. F., Helmkampf, M. & Bornberg-Bauer, E. Mechanisms and dynamics of orphan gene emergence in insect genomes. Genome Biol. Evol. 5, 439–455 (2013).

13. Thomas, G. W. C. et al. Gene content evolution in the arthropods. Genome Biol. 21, 15 (2020).

14. Dou, X. Y., Zhang, A. J. & Jurenka, R. Functional identification of fatty acyl reductases in female pheromone gland and tarsi of the corn earworm, *Helicoverpa zea*. Insect Biochem. Mol. Biol. 116, 103260 (2020).

15. Carot-Sans, G., Muñoz, L., Piulachs, M. D., Guerrero, A. & Rosell, G. Identification and characterization of a fatty acyl reductase from a *Spodoptera littoralis* female gland involved in pheromone biosynthesis. Insect Mol. Biol. 24, 82–92 (2015).

16. Teerawanichpan, P., Robertson, A. J. & Qiu, X. A fatty acyl-CoA reductase highly expressed in the head of honey bee (*Apis mellifera*) involves biosynthesis of a wide range of aliphatic fatty alcohols. Insect Biochem. Mol. Biol. 40, 641–649 (2010).

17. Xiong, G. et al. Body shape and coloration of silkworm larvae are influenced by a novel cuticular protein. Genetics 207, 1053–1066 (2017).

18. Pan, P. L. et al. A comprehensive omics analysis and functional survey of cuticular proteins in the brown planthopper. Proc. Natl. Acad. Sci. USA. 115, 5175–5180 (2018).

19. Xu, H. X. et al. Comparative genomics sheds light on the convergent evolution of miniaturized wasps. Mol. Biol. Evol. 38, 5539–5554 (2021).

20. Zhao, X. M., Su, L., Xu, W. L., Schaack, S. & Sun, C. Genome-wide identification of accessible chromatin regions in bumblebee by ATAC-seq. Sci. Data 7, 367 (2020).

21. Kapheim, K. M. et al. Genomic signatures of evolutionary transitions from solitary to group living. Science 348, 1139–1143 (2015).

22. Woodard, S. H. et al. Genes involved in convergent evolution of eusociality in bees. Proc. Natl. Acad. Sci. USA 108, 7472–7477 (2011).

23. Lacy, K. D. & Kronauer, D. J. C. Evolution: how sweat bees gained and lost eusociality. Curr. Biol. 33, R770–R773 (2023).

24. Mikhailova, A. A., Rinke, S. & Harrison, M. C. Genomic signatures of eusocial evolution in insects. Curr. Opin. Insect Sci. 61, 101136 (2024).

25. Jones, B. M. et al. Convergent and complementary selection shaped gains and losses of eusociality in sweat bees. *Nat*. Ecol. Evol. 7, 557–569 (2023).

26. Wyatt, C. D. R. et al. Social complexity, life-history and lineage influence the molecular basis of castes in vespid wasps. Nat. Commun. 14, 1046 (2023).

27. Vilhelmsen, L. Larval anatomy of Orussidae (Hymenoptera). J. Hymenopt. Res. 12, 346–354 (2003).

28. Chen, Y. Y. et al. Female semiochemicals stimulate male courtship but dampen female sexual receptivity. Proc. Natl. Acad. Sci. USA 120, e2311166120.

29. Chertemps, T. et al. A female-biased expressed elongase involved in long-chain hydrocarbon biosynthesis and courtship behavior in *Drosophila melanogaster*. Proc. Natl. Acad. Sci. USA 104, 4273–4278 (2007).

30. Rim, E. Y., Clevers, H. & Nusse, R. The Wnt pathway: from signaling mechanisms to synthetic modulators. Annu. Rev. Biochem. 91, 571–598 (2022).

31. Zhou, B. H. et al. Notch signaling pathway: architecture, disease, and therapeutics. Signal Transduct. Target. Ther. 7, 95 (2022).

32. Richards, E. H. Salivary secretions from the ectoparasitic wasp, *Eulophus pennicornis* contain hydrolases, and kill host hemocytes by apoptosis. Arch. Insect Biochem. Physiol. 79, 61–74 (2012).

33. Shi, J. M. et al. The larval saliva of an endoparasitic wasp, *Pteromalus puparum*, suppresses host immunity. J. Insect Physiol. 141, 104425 (2022).

34. Pang, L. et al. Larval secretions of parasitoid wasps are new effectors that impair host immune defences. Crop Health 1, 11 (2023).

35. Peterson, C. L. & Laniel, M.-A. Histones and histone modifications. Curr. Biol. 14, R546–R551 (2004).

36. Martire, S. & Banaszynski, L. A. The roles of histone variants in fine-tuning chromatin organization and function. Nat. Rev. Mol. Cell Biol. 21, 522–541 (2020).

37. Kurata, S. Peptidoglycan recognition proteins in *Drosophila* immunity. Dev. Comp. Immunol. 42, (2014).

38. Mei, X. H. et al. Peptidoglycan recognition protein 6 (PGRP6) from Asian corn borer, *Ostrinia furnacalis* (Guenée) serve as a pattern recognition receptor in innate immune response. Arch. Insect Biochem. Physiol. 111, e21955 (2022).

39. Xiao, D. Y. et al. The roles of SMYD4 in epigenetic regulation of cardiac development in zebrafish. PLoS Genet. 14, (2018).

40. International Helminth Genomes Consortium. Comparative genomics of the major parasitic worms. Nat. Genet. 51, 163–174 (2019).

41. Lu, T. M., Kanda, M., Furuya, H. & Satoh, N. Dicyemid mesozoans: a unique parasitic lifestyle and a reduced genome. Genome Biol. Evol. 11, 2232–2243 (2019).

42. Ralton, J. E., Sernee, M. F. & McConville, M. J. Evolution and function of carbohydrate reserve biosynthesis in parasitic protists. Trends Parasitol. 37, 988–1001 (2021).

43. Chen, X. L. et al. *Balanophora* genomes display massively convergent evolution with other extreme holoparasites and provide novel insights into parasite–host interactions. Nat. Plants 9, 1627–1642 (2023).

44. Asgari, S. & Rivers, D. B. Venom proteins from endoparasitoid wasps and their role in host-parasite interactions. Annu. Rev. Entomol. 56, 313–335 (2011).

45. Moreau, S. & Asgari, S. Venom proteins from parasitoid wasps and their biological functions. Toxins (Basel) 7, 2385–2412 (2015).

46. Teng, Z. W. et al. Protein discovery: combined transcriptomic and proteomic analyses of venom from the endoparasitoid *Cotesia chilonis* (Hymenoptera: Braconidae). Toxins (Basel) 9, 135 (2017).

47. Zhao, W. et al. Comparative transcriptome analysis of venom glands from *Cotesia vestalis* and *Diadromus collaris*, two endoparasitoids of the host *Plutella xylostella*. Sci. Rep. 7, (2017).

48. Goecks, J. et al. Integrative approach reveals composition of endoparasitoid wasp venoms. PLoS One 8, e64125 (2013).

49. Burke, G. R. & Strand, M. R. Systematic analysis of a wasp parasitism arsenal. Mol. Ecol. 23, 890–901 (2014).

50. Werren, J. H. et al. Functional and evolutionary insights from the genomes of three parasitoid *Nasonia* species. Science 327, 343–348 (2010).

51. Dennis, A. B. et al. Functional insights from the GC-poor genomes of two aphid parasitoids, Aphidius ervi and Lysiphlebus fabarum. BMC Genom. 21, 376 (2020).

52. Formesyn, E. M., Heyninck, K. & de Graaf, D. C. The role of serine- and metalloproteases in *Nasonia vitripennis* venom in cell death related processes towards a *Spodoptera frugiperda* Sf21 cell line. J. Insect Physiol. 59, 795–803 (2013).

53. Ye, X. H. et al. Comprehensive isoform-level analysis reveals the contribution of alternative isoforms to venom evolution and repertoire diversity. Genome Res. 33, 1554–1567 (2023).

54. Yang, L. et al. Identification and comparative analysis of venom proteins in a pupal ectoparasitoid, *Pachycrepoideus vindemmiae*. Front. Physiol. 11, 9 (2020).

55. Lin, Z. et al. Insights into the venom protein components of *Microplitis mediator*, an endoparasitoid wasp. Insect Biochem. Mol. Biol. 105, 33–42 (2019).

56. Parkinson, N., Conyers, C. & Smith, I. A venom protein from the endoparasitoid wasp *Pimpla hypochondriaca* is similar to snake venom reprolysin-type metalloproteases. J. Invertebr. Pathol. 79, 129–131 (2002).

57. Price, D. R. G. et al. A venom metalloproteinase from the parasitic wasp *Eulophus pennicornis* is toxic towards its host, tomato moth (*Lacanobia oleracae*). Insect Mol. Biol. 18, 195–202 (2009).

58. Dorémus, T. et al. Venom gland extract is not required for successful parasitism in the polydnavirus-associated endoparasitoid *Hyposoter didymator* (Hym. Ichneumonidae) despite the presence of numerous novel and conserved venom proteins. Insect Biochem. Mol. Biol. 43, 292–307 (2013).

59. Alvarado, G. et al. Bioinformatic analysis suggests potential mechanisms underlying parasitoid venom evolution and function. Genomics 112, 1096–1104 (2020).

60. Vincent, B. et al. The venom composition of the parasitic wasp *Chelonus inanitus* resolved by combined expressed sequence tags analysis and proteomic approach. BMC Genom. 11, 693 (2010).

61. Gatti, J.-L. et al. Proteo-trancriptomic analyses reveal a large expansion of metalloprotease-like proteins in atypical venom vesicles of the wasp *Meteorus pulchricornis* (Braconidae). Toxins (Basel) 13, 502 (2021).

62. Yu, K. L. et al. Multi-omic identification of venom proteins collected from artificial hosts of a parasitoid wasp. Toxins (Basel) 15, 377 (2023).

63. Lin, Z. et al. A metalloprotease homolog venom protein from a parasitoid wasp suppresses the Toll pathway in host hemocytes. Front. Immunol. 9, 2301 (2018).

64. Yang, Y. et al. Genome of the pincer wasp *Gonatopus flavifemur* reveals unique venom evolution and a dual adaptation to parasitism and predation. BMC Biol. 19, 145 (2021).

65. Colinet, D. et al. Extensive inter- and intraspecific venom variation in closely related parasites targeting the same host: the case of *Leptopilina* parasitoids of *Drosophila*. Insect Biochem. Mol. Biol. 43, 601–611 (2013).

66. Crawford, A. M. et al. The constituents of *Microctonus sp.* parasitoid venoms. Insect Mol. Biol. 17, 313–324 (2008).

67. Zhu, J. Y. Deciphering the main venom components of the ectoparasitic ant-like bethylid wasp, *Scleroderma guani*. Toxicon 113, 32–40 (2016).

68. Liu, N. Y. et al. Venomics reveals novel ion transport peptide-likes (ITPLs) from the parasitoid wasp *Tetrastichus brontispae*. Toxicon 141, 88–93 (2018).

69. Becchimanzi, A. et al. Venomics of the ectoparasitoid wasp *Bracon nigricans*. BMC Genom. 21, 34 (2020).

70. Ye, X. H. et al. Genomic signatures associated with maintenance of genome stability and venom turnover in two parasitoid wasps. Nat. Commun. 13, 6417 (2022).

71. Colinet, D. et al. Identification of the main venom protein components of *Aphidius ervi*, a parasitoid wasp of the aphid model *Acyrthosiphon pisum*. BMC Genom. 15, 342 (2014).

72. Zhou, L. Z. et al. Two venom serpins from the parasitoid wasp *Microplitis mediator* inhibit the host prophenoloxidase activation and antimicrobial peptide synthesis. Insect Biochem. Mol. Biol. 152, 103895 (2023).

73. Yan, Z. C. et al. A serpin gene from a parasitoid wasp disrupts host immunity and exhibits adaptive alternative splicing. PLoS Pathog. 19, e1011649 (2023).

74. Zhao, X. X., et al. Genomic signatures associated with the evolutionary loss of egg yolk in parasitoid wasps. Preprint at 10.1101/2023.12.30.573744 (2024).

75. Ralec, A. Egg contents in relation to host-feeding in some parasitic hymenoptera. Entomophaga 40, 87–93 (1995).

76. Grbic, M. & Strand, M. R. Shifts in the life history of parasitic wasps correlate with pronounced alterations in early development. Proc. Natl. Acad. Sci. USA 95, 1097–1101 (1998).

77. Donnell, D. M. Vitellogenin of the parasitoid wasp, *Encarsia formosa* (Hymenoptera: Aphelinidae): gene organization and differential use by members of the genus. Insect Biochem. Mol. Biol. 34, 951–961 (2004).

78. Hegedus, D., Erlandson, M., Gillott, C. & Toprak, U. New insights into peritrophic matrix synthesis, architecture, and function. Annu. Rev. Entomol. 54, 285–302 (2009).

79. Zhu, L. F. et al. Adaptive evolution to a high purine and fat diet of carnivorans revealed by gut microbiomes and host genomes. Environ. Microbiol. 20, 1711–1722 (2018).

80. Gloss, A. D. et al. Evolution in an ancient detoxification pathway is coupled with a transition to herbivory in the Drosophilidae. Mol. Biol. Evol. 31, 2441 (2014).

81. Gloss, A. D. et al. Evolution of herbivory remodels a *Drosophila* genome. Preprint at 10.1101/767160 (2019).

82. Winde, I. & Wittstock, U. Insect herbivore counteradaptations to the plant glucosinolate–myrosinase system. Phytochemistry 72, 1566–1575 (2011).

83. Lv, Q. Q., Li, X. F., Fan, B. F., Zhu, C. & Chen, Z. X. The cellular and subcellular organization of the glucosinolate–myrosinase system against herbivores and pathogens. Int. J. Mol. Sci. 23, 1577 (2022).

84. Abdalsamee, M. K., Giampà, M., Niehaus, K. & Müller, C. Rapid incorporation of glucosinolates as a strategy used by a herbivore to prevent activation by myrosinases. Insect Biochem. Mol. Biol. 52, 115–123 (2014).

85. Müller, C. Interactions between glucosinolate- and myrosinase-containing plants and the sawfly *Athalia rosae*. Phytochem. Rev. 8, 121–134 (2009).

86. Bridges, M. et al. Spatial organization of the glucosinolate-myrosinase system in brassica specialist aphids is similar to that of the host plant. Proc. Biol. Sci. 269, 187–191 (2002).

87. Gurevich, A., Saveliev, V., Vyahhi, N. & Tesler, G. QUAST: quality assessment tool for genome assemblies. Bioinformatics 29, 1072–1075 (2013).

88. Manni, M., Berkeley, M. R., Seppey, M. & Zdobnov, E. M. BUSCO: assessing genomic data quality and beyond. Curr. Protoc. 1, e323 (2021).

89. Derelle, R., Philippe, H. & Colbourne, J. K. Broccoli: combining phylogenetic and network analyses for orthology assignment. Mol. Biol. Evol. 37, 3389–3396 (2020).

90. Katoh, K., Misawa, K., Kuma, K. & Miyata, T. MAFFT: a novel method for rapid multiple sequence alignment based on fast Fourier transform. Nucleic Acids Res. 30, 3059–3066 (2002).

91. Emms, D. M. & Kelly, S. OrthoFinder: phylogenetic orthology inference for comparative genomics. Genome Biol. 20, 238 (2019).

92. Minh, B. Q. et al. IQ-TREE 2: new models and efficient methods for phylogenetic inference in the genomic era. Mol. Biol. Evol. 37, 1530–1534 (2020).

93. Kalyaanamoorthy, S., Minh, B. Q., Wong, T. K., von Haeseler, A. & Jermiin, L. S. ModelFinder: fast model selection for accurate phylogenetic estimates. Nat. Methods 14, 587–589 (2017).

94. Sanderson, M. J. r8s: inferring absolute rates of molecular evolution and divergence times in the absence of a molecular clock. Bioinformatics 19, 301–302 (2003).

95. Mendes, F. K., Vanderpool, D., Fulton, B. & Hahn, M. W. CAFE 5 models variation in evolutionary rates among gene families. Bioinformatics 36, 5516–5518 (2021).

96. Pond, S. L. K., Frost, S. D. W. & Muse, S. V. HyPhy: hypothesis testing using phylogenies. Bioinformatics 21, 676–679 (2005).

97. Smith, M. D. et al. Less is more: an adaptive branch-site random effects model for efficient detection of episodic diversifying selection. Mol. Biol. Evol. 32, 1342–1353 (2015).

98. Yang, Z. H. PAML 4: phylogenetic analysis by maximum likelihood. Mol. Biol. Evol. 24, 1586–1591 (2007).

99. Dohmen, E., Klasberg, S., Bornberg-Bauer, E., Perrey, S. & Kemena, C. The modular nature of protein evolution: domain rearrangement rates across eukaryotic life. BMC Evol. Biol. 20, 30 (2020).

100. Cantalapiedra, C. P., Hernández-Plaza, A., Letunic, I., Bork, P. & Huerta-Cepas, J. eggNOG-mapper v2: functional annotation, orthology assignments, and domain prediction at the metagenomic scale. Mol. Biol. Evol. 38, 5825–5829 (2021).

101. Klopfenstein, D. V. et al. GOATOOLS: a python library for gene ontology analyses. Sci. Rep. 8, 10872 (2018).

102. Armstrong, J. et al. Progressive Cactus is a multiple-genome aligner for the thousand-genome era. Nature 587, 246–251 (2020).

103. Hubisz, M. J., Pollard, K. S. & Siepel, A. PHAST and RPHAST: phylogenetic analysis with space/time models. Brief. Bioinform. 12, 41–51 (2011).

104. Siepel, A. et al. Evolutionarily conserved elements in vertebrate, insect, worm, and yeast genomes. Genome Res. 15, 1034–1050 (2005).

105. Bolger, A. M., Lohse, M. & Usadel, B. Trimmomatic: a flexible trimmer for Illumina sequence data. Bioinformatics 30, 2114–2120 (2014).

106. Langmead, B. & Salzberg, S. L. Fast gapped-read alignment with Bowtie 2. Nat. Methods 9, 357–359 (2012).

107. Li, H. et al. The sequence alignment/map format and SAMtools. Bioinformatics 25, 2078–2079 (2009).

108. Gaspar, J. M. Improved peak-calling with MACS2. Preprint at 10.1101/496521 (2018).

109. Chen, S. F., Zhou, Y. Q., Chen, Y. R. & Gu, J. fastp: an ultra-fast all-in-one FASTQ preprocessor. Bioinformatics 34, i884–i890 (2018).

110. Patro, R., Duggal, G., Love, M. I., Irizarry, R. A. & Kingsford, C. Salmon provides fast and bias-aware quantification of transcript expression. Nat. Methods 14, 417–419 (2017).

