## Supplementary Figures for "Large-scale genome analyses provide insights into Hymenoptera evolution"

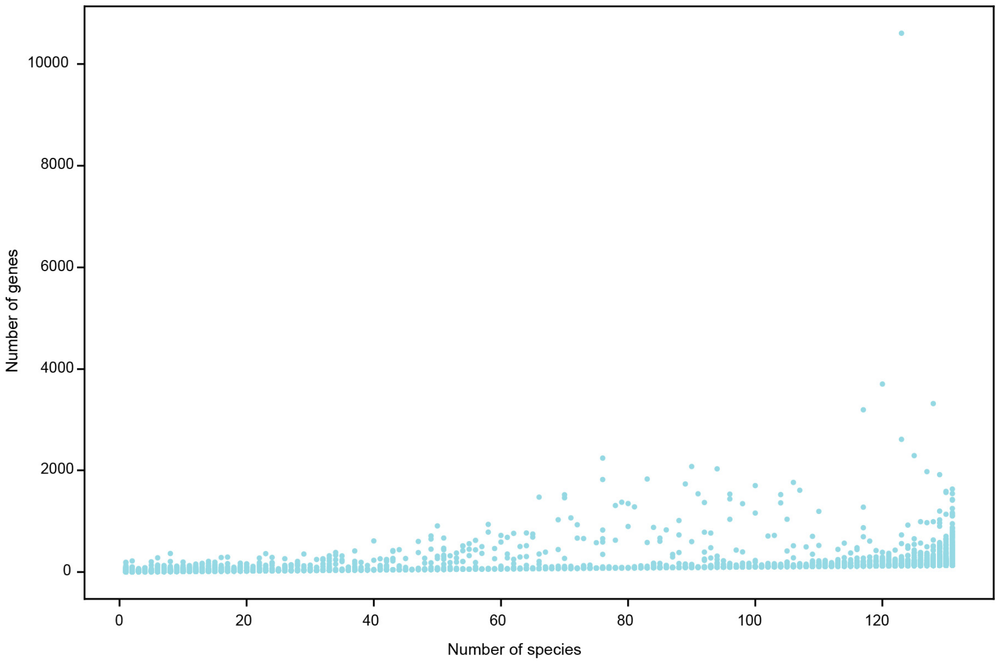


**Supplementary Fig. 1 | A scatter plot of gene family size against the number of species that include the family.** Every point indicates a gene family.

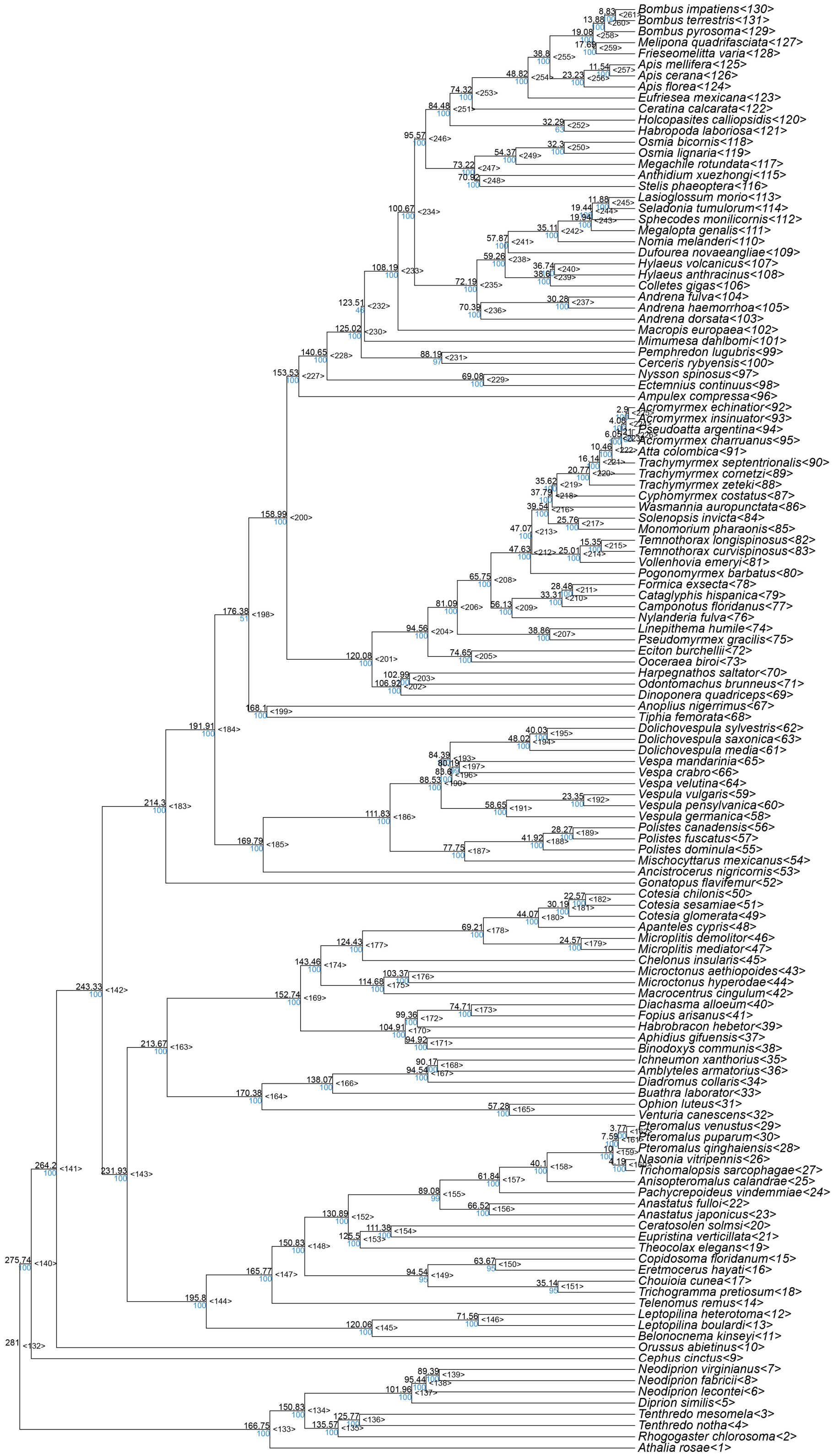


**Supplementary Fig. 2 | Phylogenetic relationships among 131 Hymenoptera insects.** The phylogenetic tree is identical to the one shown in Fig. 1, but additionally shows the estimated divergence times in million years ago (black numbers, upper left of nodes), bootstrap support (blue numbers, lower left of nodes), and node ID (in parentheses next to the node).

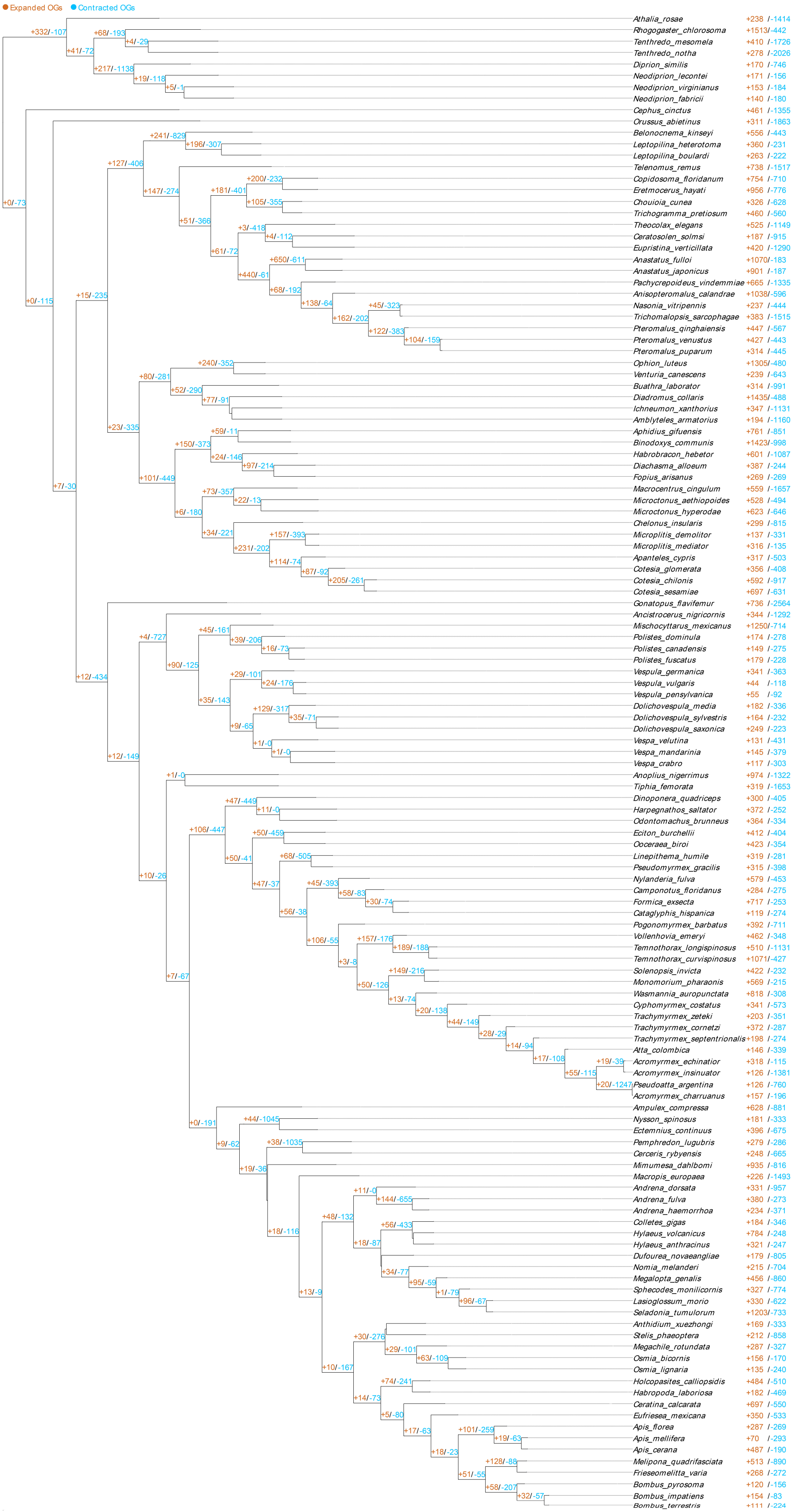


**Supplementary Fig. 3 | Gene family expansion and contraction throughout 131 hymenopteran insects.** Yellow numbers indicate expanded gene families, and blue numbers indicate contracted gene families.

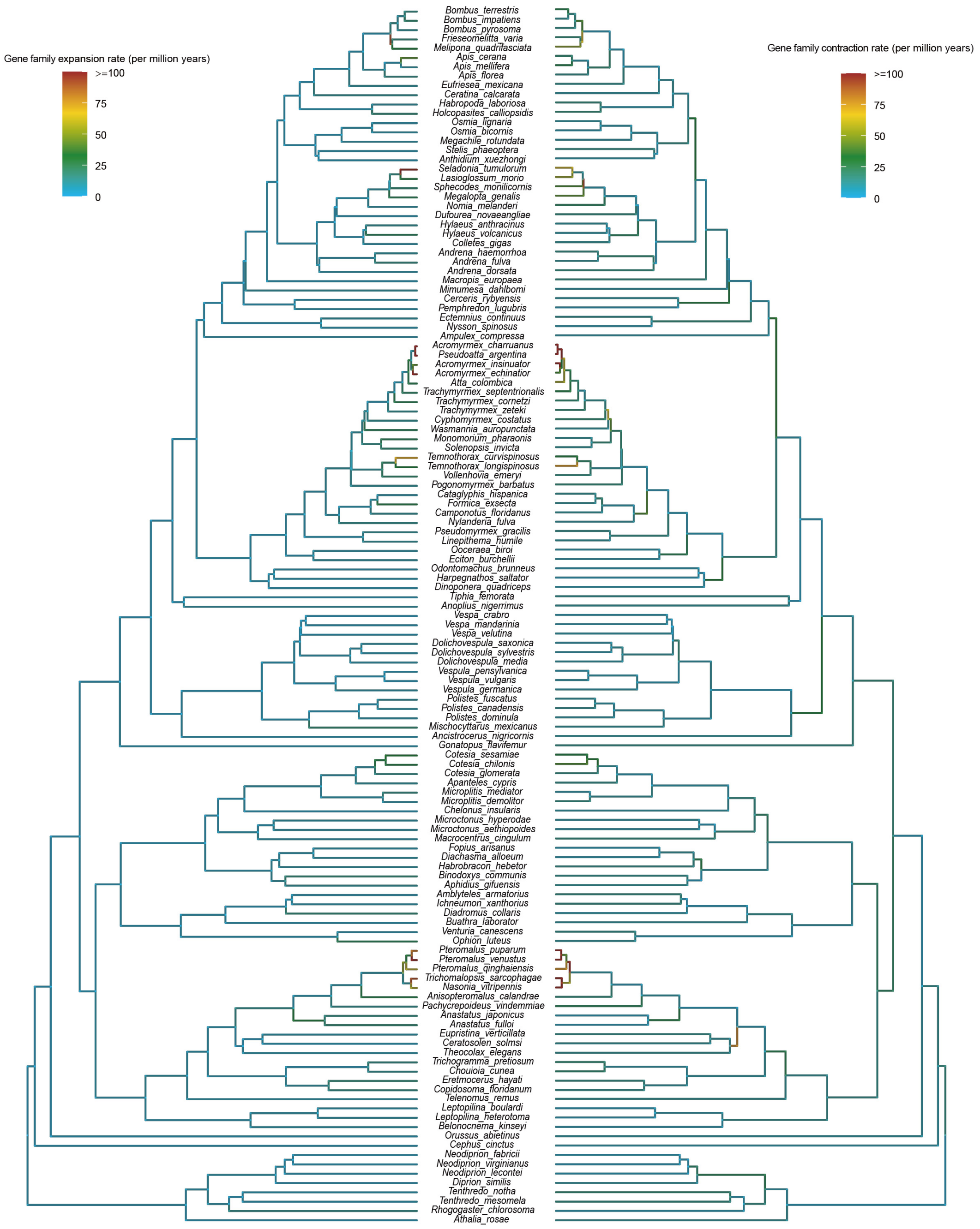


**Supplementary Fig. 4 | Gene family expansion and contraction rate throughout all species.** The gene family expansion and contraction rate refer to the number of gene families that expanded or contracted per million years on a particular branch, respectively.

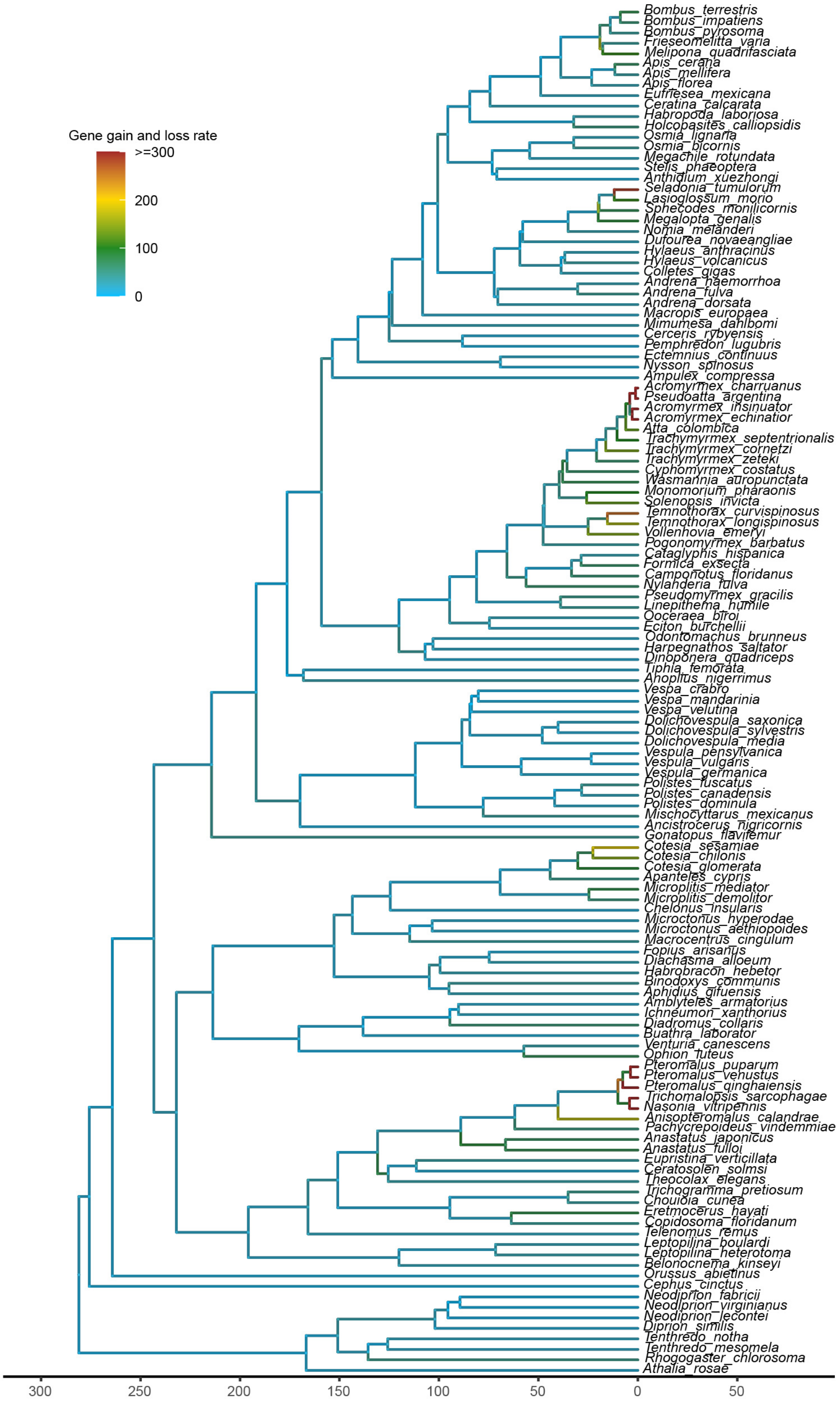


**Supplementary Fig. 5 | Gene gain and loss rate throughout all species.** The gene gain and loss rate refers to the sum of gene gains and losses per million years on a particular branch. The clade comprising *Nasonia*, *Trichomalopsis*, and *Pteromolaus* in Pteromalidae and branches of leafcutter ants (Formicidae) exhibit noticeable accelerations in gene gain and loss rates.

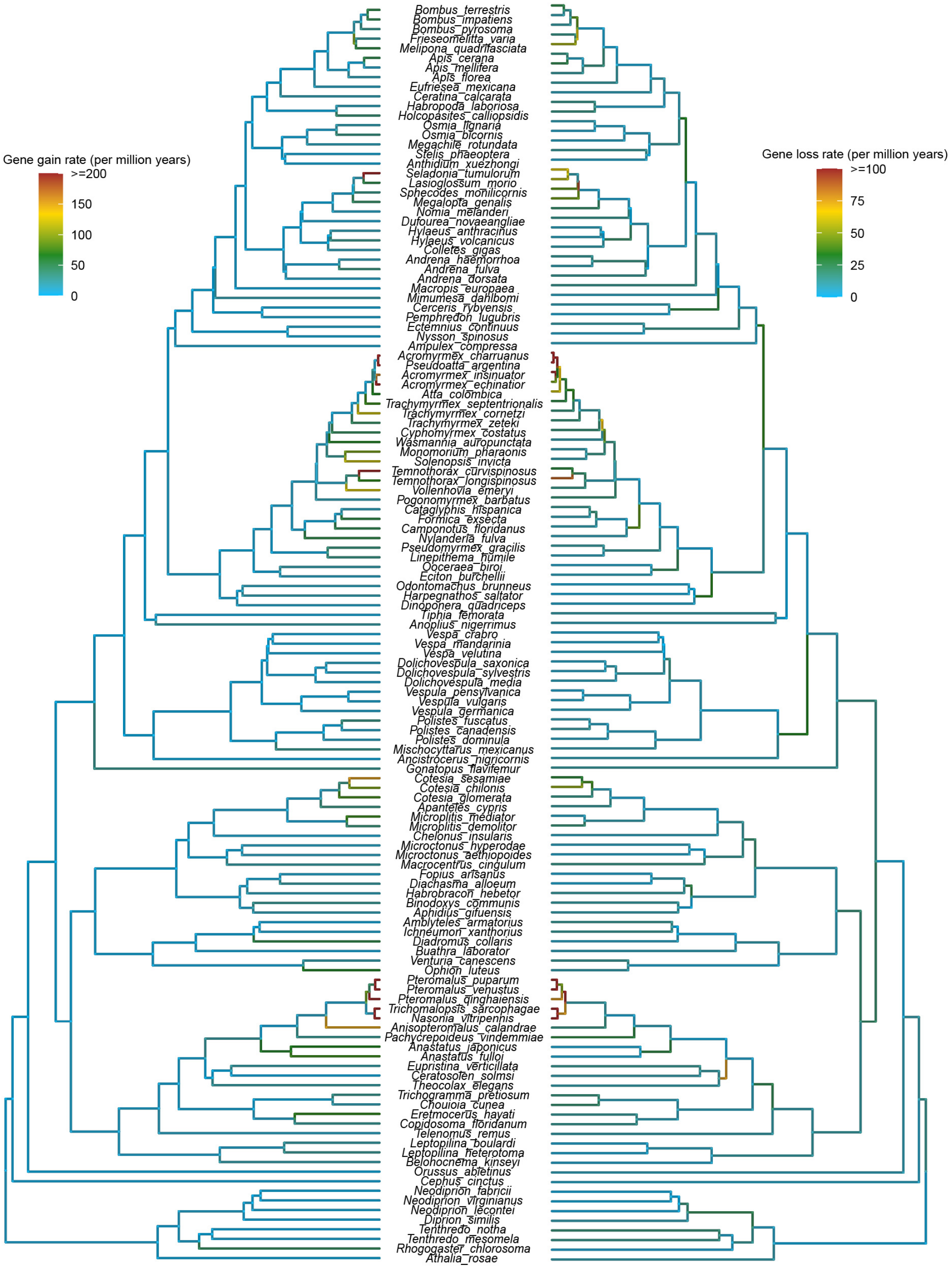


**Supplementary Fig. 6 | Gene gain and loss rates varied across species.** Gene gain and loss rate refers to the gene gains and losses per million years on a particular branch, respectively.

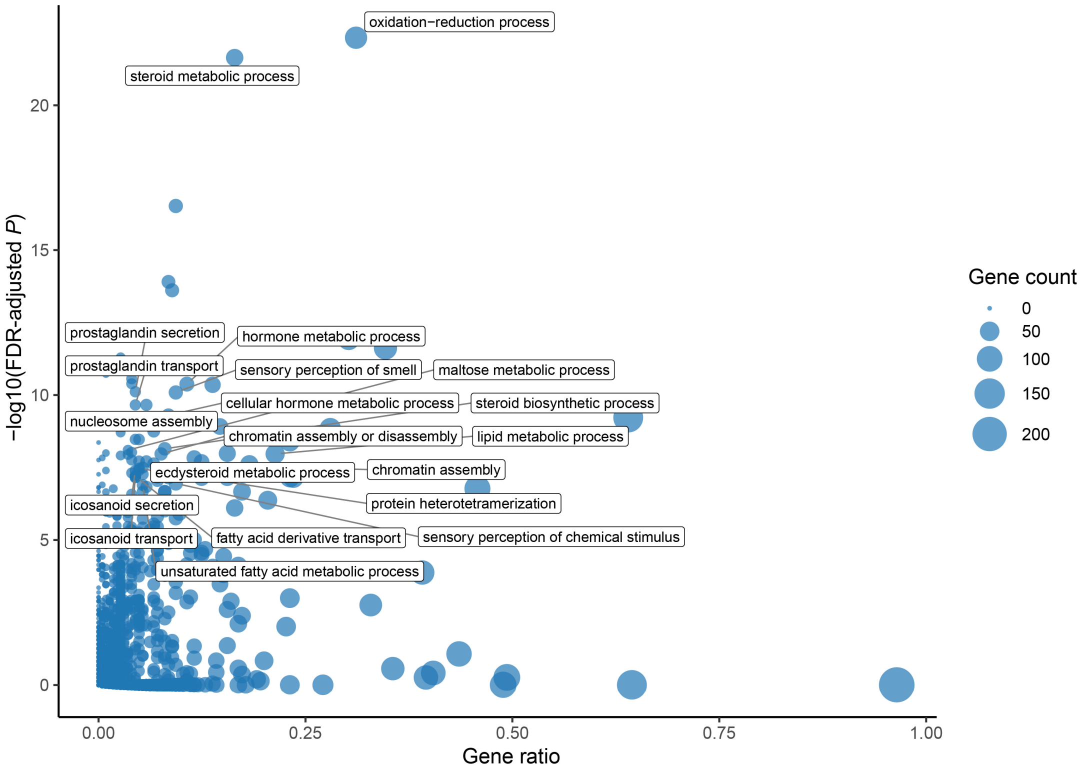


**Supplementary Fig. 7 | GO enrichment analysis of gene families that exhibited significant expansions or contractions during Hymenoptera evolution (CAFE5 *P* value < 0.05).** The gray box shows the GO terms for the 20 biological processes with the lowest adjusted *P* value (FDR-adjusted *P* < 0.05, Benjamini-Hochberg multi-test correction).

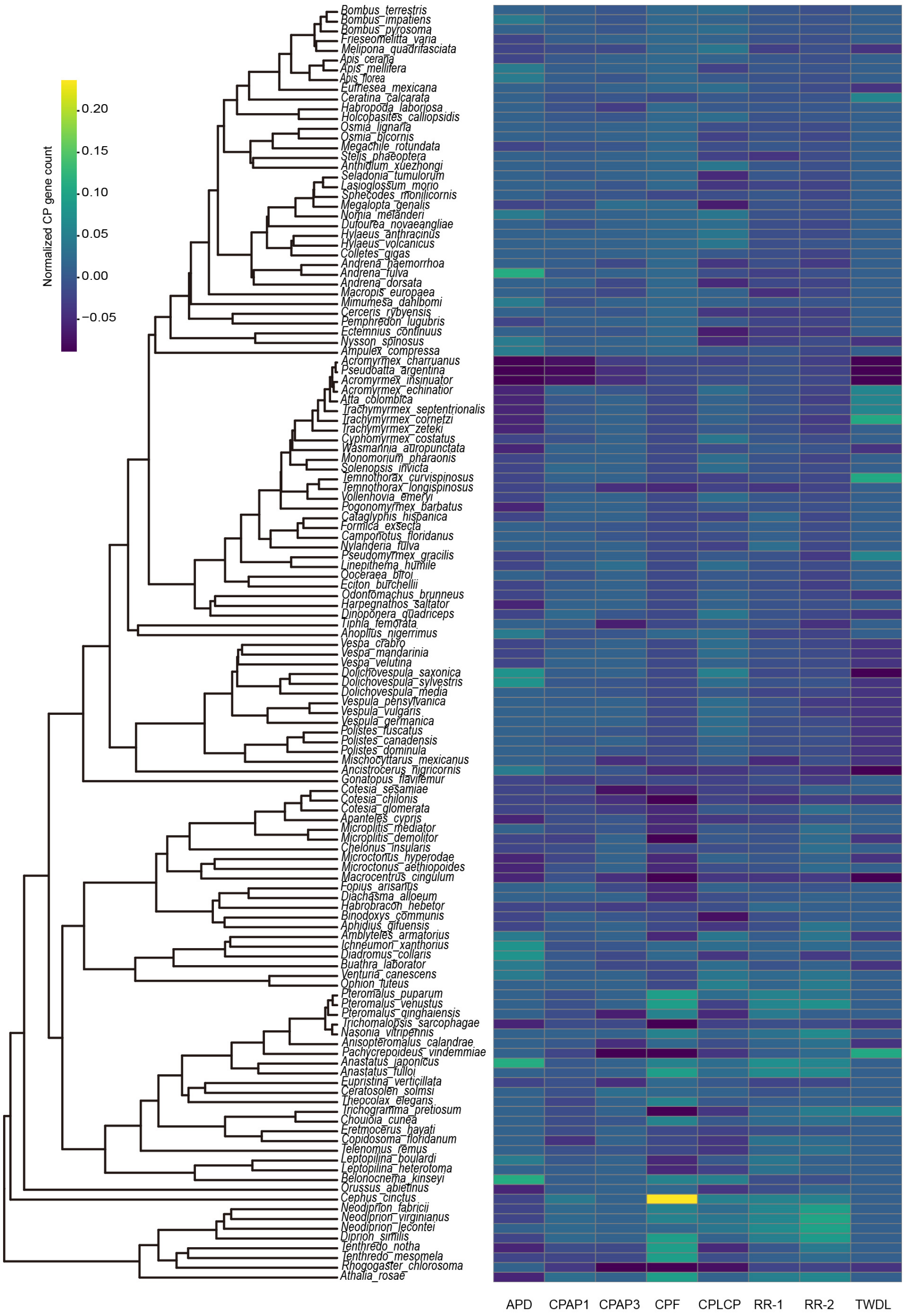


**Supplementary Fig. 8 | The size distribution of cuticular protein families throughout all species.** Gene numbers are normalized by the z-score. Shown from left to right are the Apidermin (APD) family, the cuticular proteins analogous to the peritrophins (CPAP) family with one or three CBM_14 domains, the CPF family, the cuticular protein of low complexity with the proline-rich (CPLCP) family, the CPR family (RR-1 and RR-2), and the Tweedle (TWDL) family.

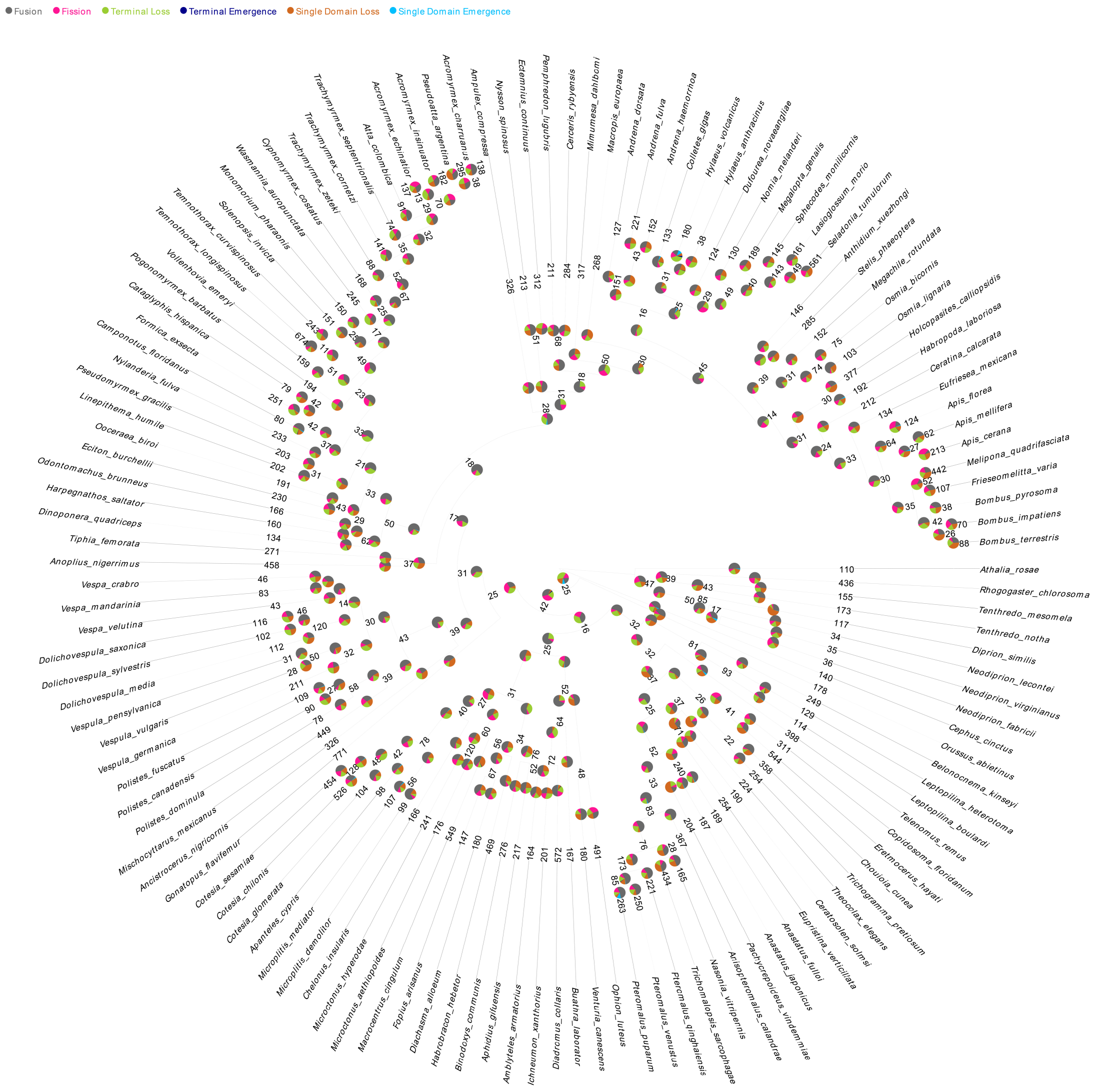


**Supplementary Fig. 9 | Protein domain rearrangement events occurred throughout the hymenopteran phylogeny.** The pie chart at each node shows the proportion of each type of rearranged event (i.e., fusion, fission, terminal loss, terminal emergence, single domain loss, and single domain emergence), and the numbers show the total number of all rearranged events.

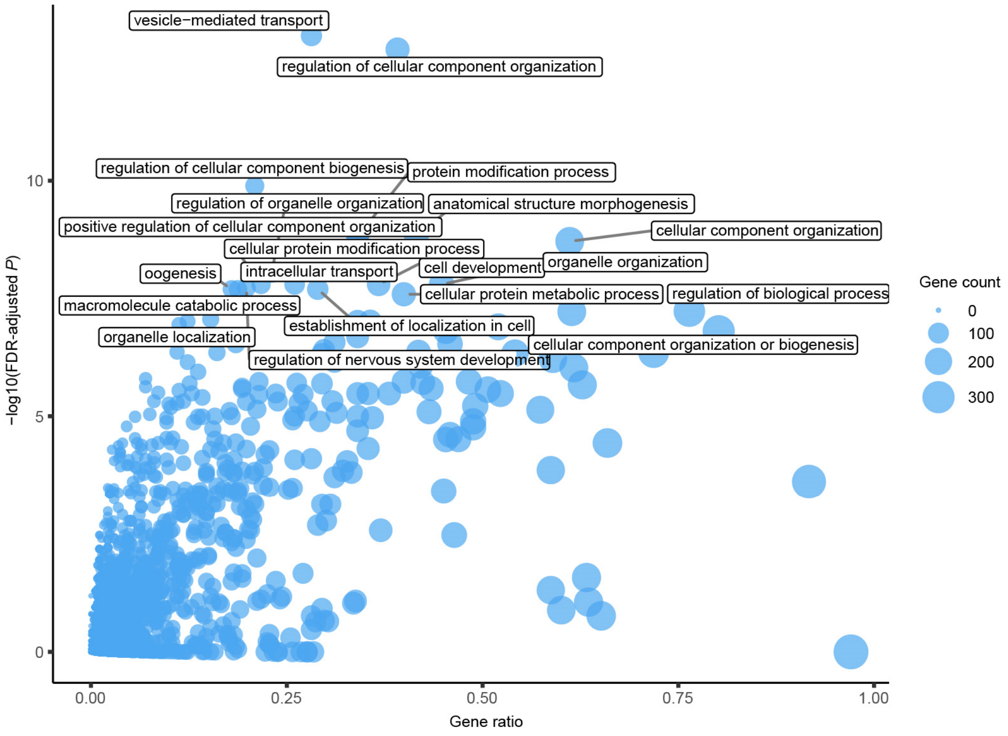


**Supplementary Fig. 10 | GO enrichment analysis of gene families with the bottom 20% evolutionary rate.** The gray box shows the GO terms for the 20 biological processes with the lowest adjusted *P* value (FDR-adjusted *P* < 0.05, Benjamini-Hochberg multi-test correction). The size of the blue circles represents the number of genes in the corresponding GO.

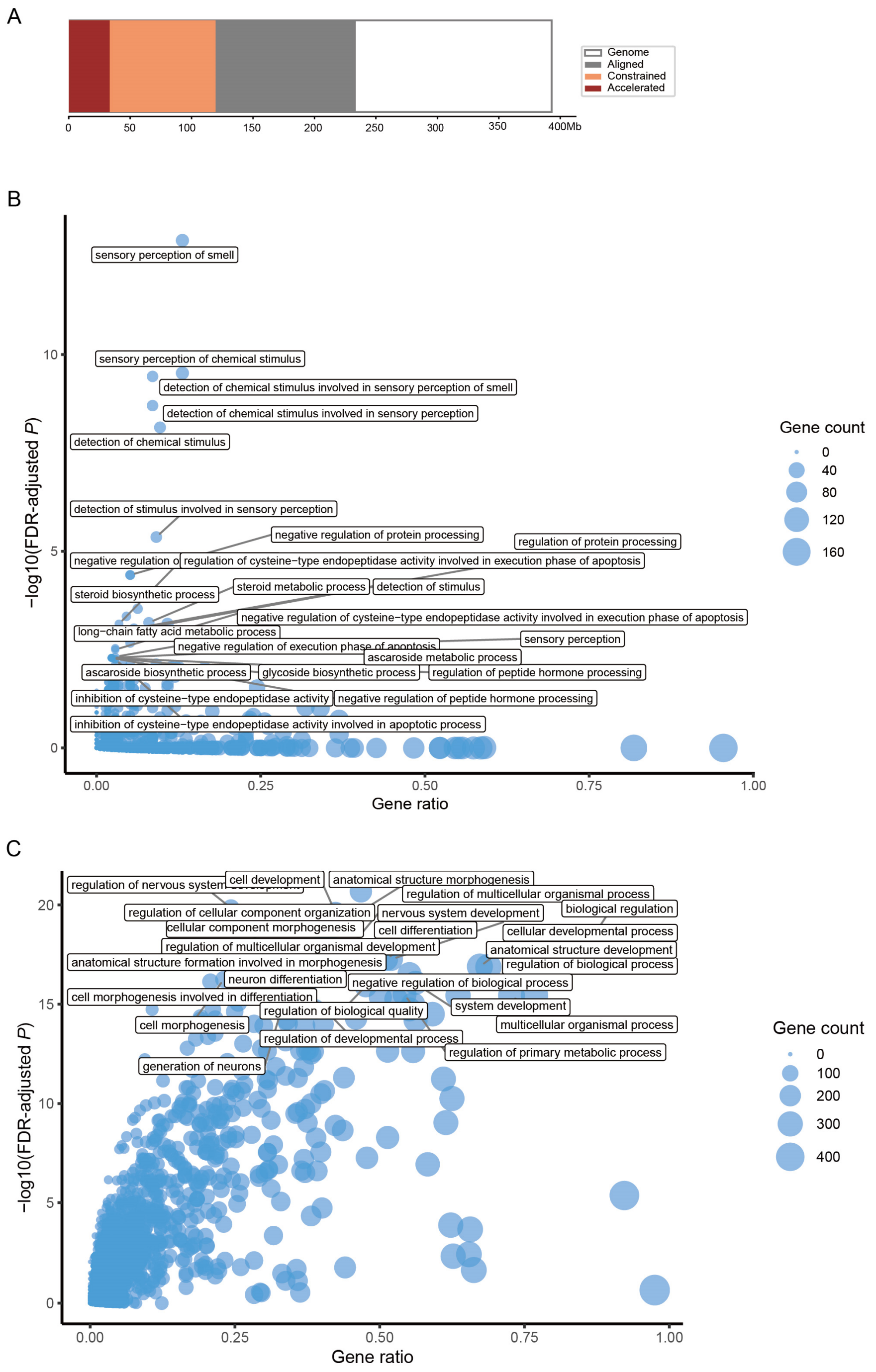


**Supplementary Fig. 11 | Analysis of nucleotides that were conserved and accelerated-evolving.** A) Nucleotide evolution analysis based on whole genome alignment. Nucleotides with negative phyloP scores are accelerated-evolving, and those with positive phyloP scores are conserved. B) GO enrichment analysis of the top 5% most accelerated genes as measured by the mean phyloP score of the coding sequence. The gray box shows the GO terms for the 25 biological processes with the lowest adjusted *P* value (FDR-adjusted *P* < 0.05, Benjamini-Hochberg multi-test correction). The size of the blue circles represents the number of genes in the corresponding GO. C) GO enrichment analysis of the top 5% most conserved genes as measured by the mean phyloP score of the coding sequence.

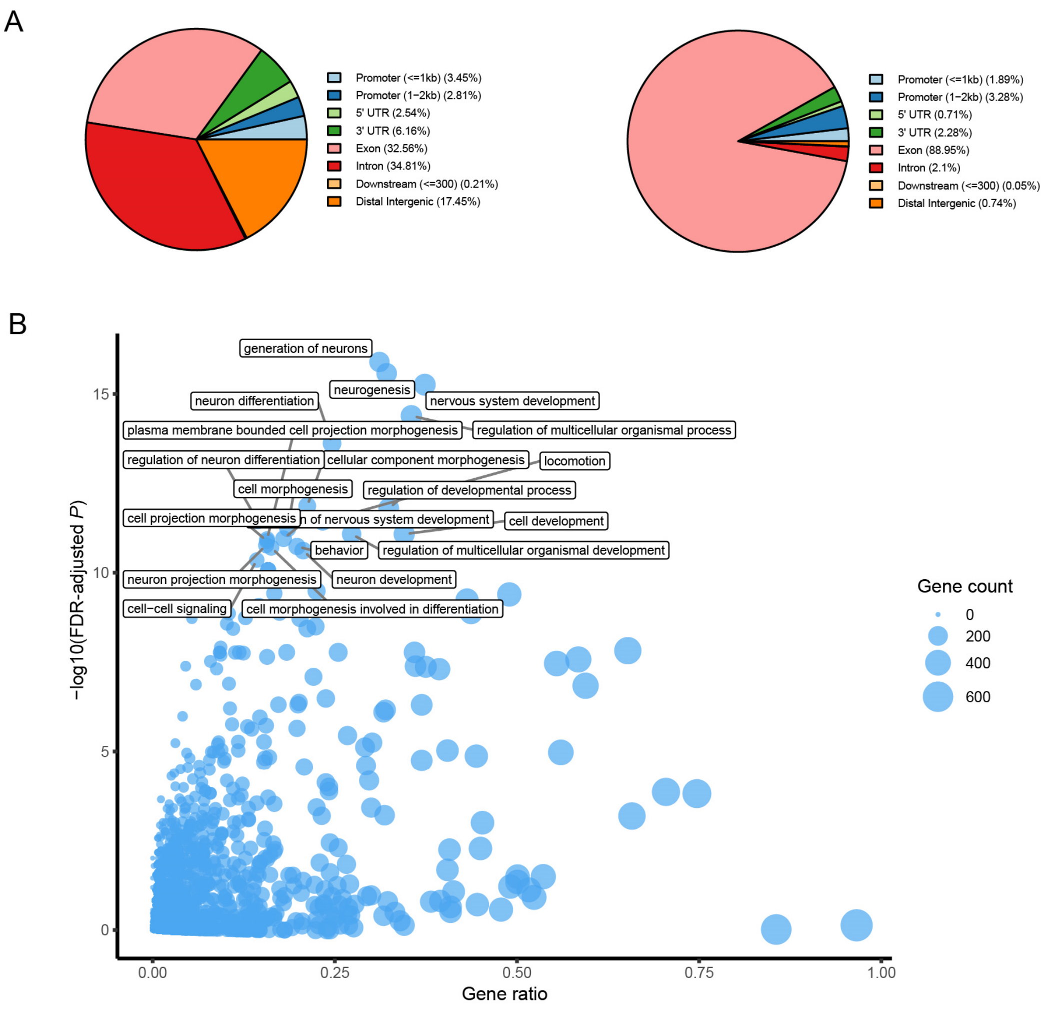


**Supplementary Fig. 12 | Annotation of conserved elements identified by phastCons and relevant gene analysis.** A) Conserved elements annotation and classification based on their relative locations to the nearest genes. The pie chart on the left is the annotation of 481,924 conserved elements, and on the right is the annotation of 27,602 elements that are conserved across all species in our 27-way alignment. B) GO enrichment analysis of genes with non-coding elements supported by ATAC-Seq within 2 kb upstream and downstream. The gray box shows the GO terms for the 20 biological processes with the lowest adjusted *P* value (FDR-adjusted $P < 0.05$, Benjamini-Hochberg multi-test correction). The size of the blue circles represents the number of genes in the corresponding GO.

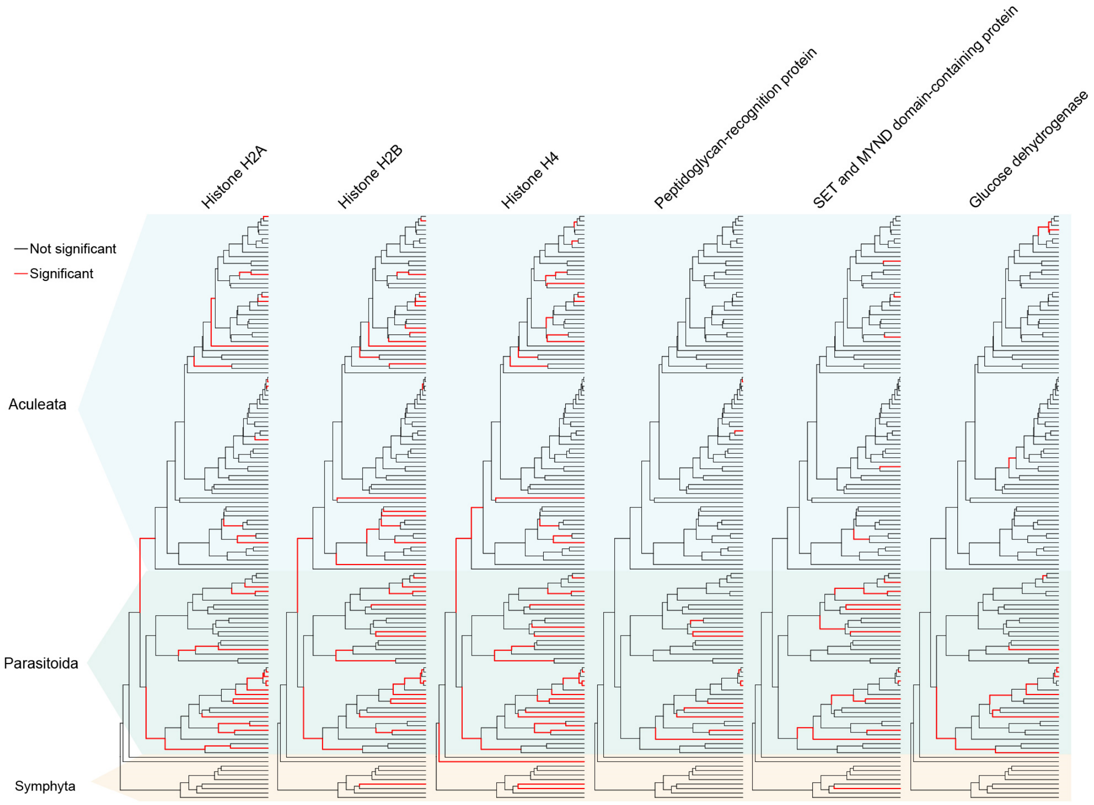


**Supplementary Fig. 13 | The distribution of the rapidly evolving events for gene families on the branches of the Parasitoida and Aculeata clades.** Six gene families, Histone H2A, Histone H2B, Histone H4, Peptidoglycan-recognition protein, SET and MYND domain-containing protein, and Glucose dehydrogenase, are shown here, which have a significantly higher occurrence rate of rapid evolutionary events in Parasitoida than in Aculeata (FDR-adjusted $P < 0.05$, Chi-square test, odds ratio >1).

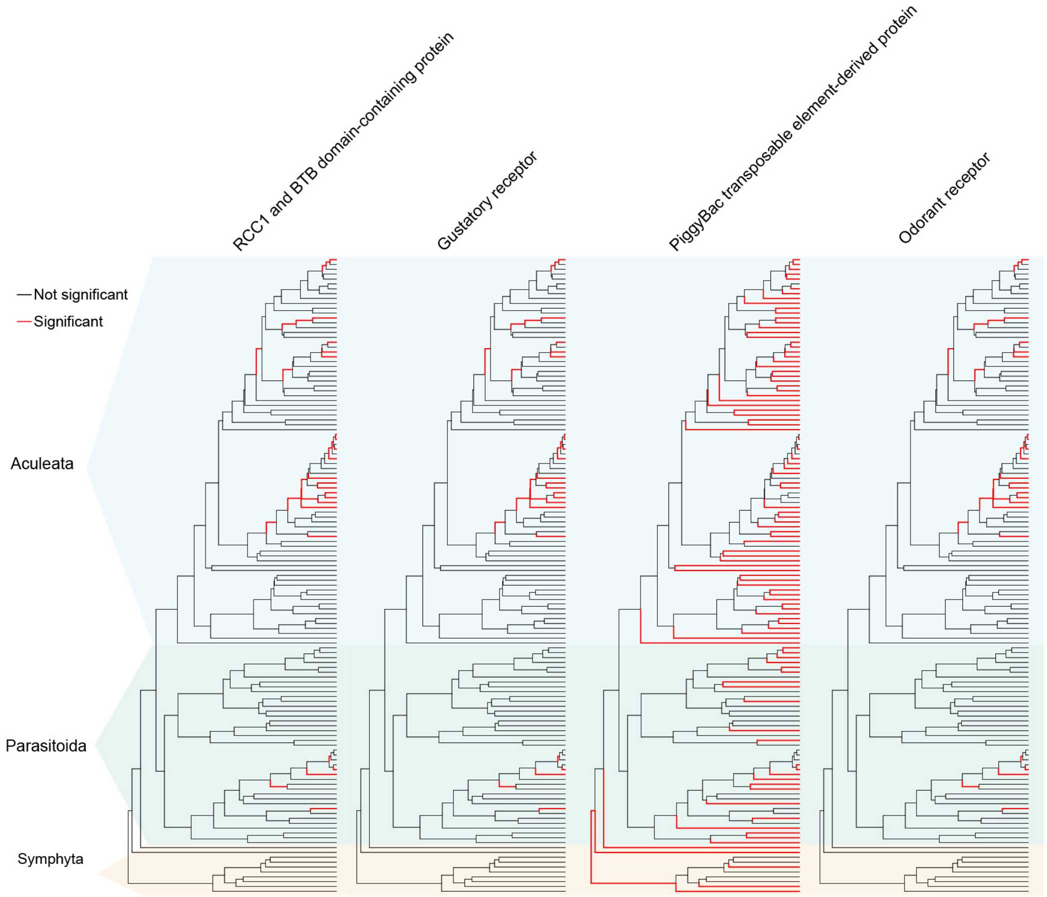


**Supplementary Fig. 14 | The distribution of the rapidly evolving events for gene families on the branches of the Parasitoida and Aculeata clades.** Four gene families, RCC1 and BTB domain-containing protein, Gustatory receptor, PiggyBac transposable element-derived protein and Odorant receptor, are shown here, which have a significantly higher occurrence rate of rapid evolutionary events in Aculeata than in Parasitoida (FDR-adjusted $P < 0.05$, Chi-square test, odds ratio >1).

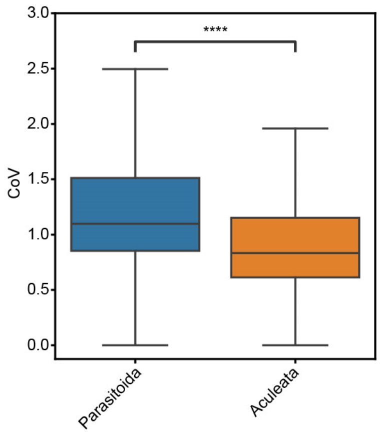


**Supplementary Fig. 15 | Comparison of the coefficient of variation (CoV) of selective constraint ($d_N/d_S$) in single-copy orthogroups between Parasitoida and Aculeata.** The one-tailed Mann-Whitney *U* test shows significantly higher CoV in selective constraint for Parasitoida compared to Aculeata (*P* = 7.92e-84).

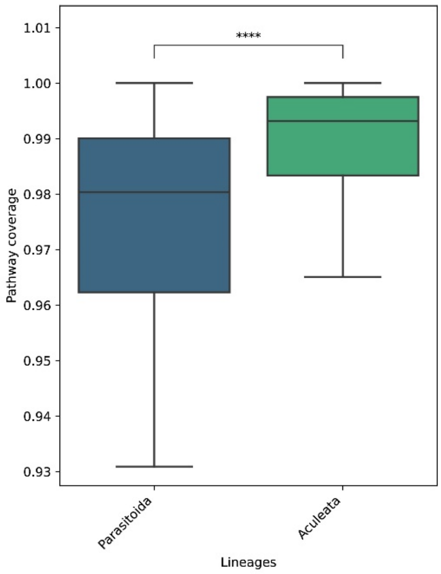


**Supplementary Fig. 16 | The overall comparison of coverage for all pathways between Parasitoida and Aculeata.** The bar plot shows significantly lower pathway coverage for Parasitoida compared to Aculeata (*P* = 2.68e-46, One-tailed Mann-Whitney *U* test). Pathway coverage is measured as the number of ECs in the pathway annotated divided by the total number of ECs in the constructed ancestral pathway.

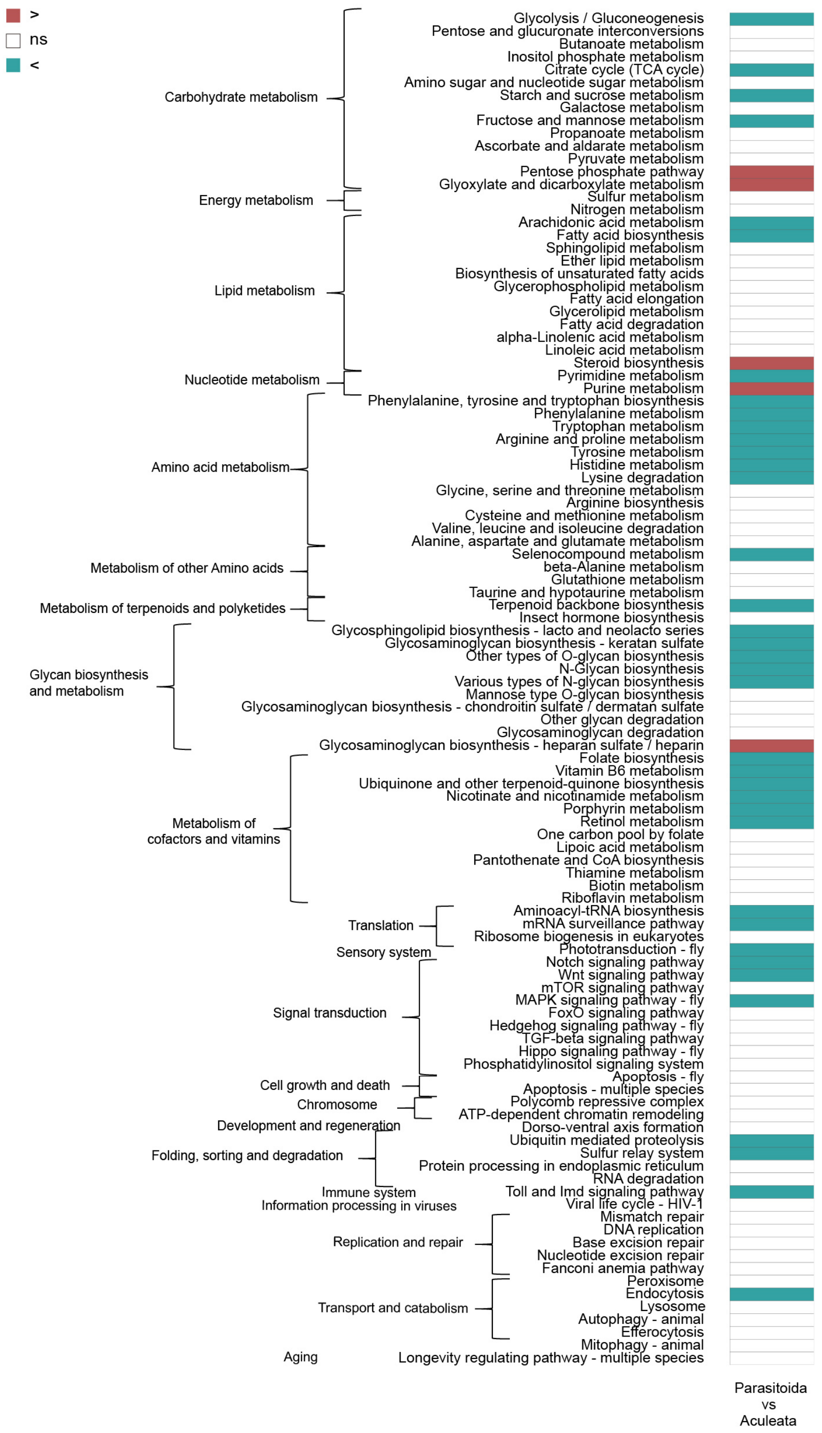


**Supplementary Fig. 17 | Pathway coverage comparison between Parasitoida and Aculeata.** Comparisons with one-sided Mann-Whitney *U* test *P*-values < 0.05 (FDR corrected using the Benjamini-Hochberg procedure) are considered significant. The color red means that the pathway coverage of the Parasitoida species is much larger than that of Aculeata; blue indicates that the Parasitoida is significantly lower than that of Aculeata; and white shows that there is no significant difference between the two clades.

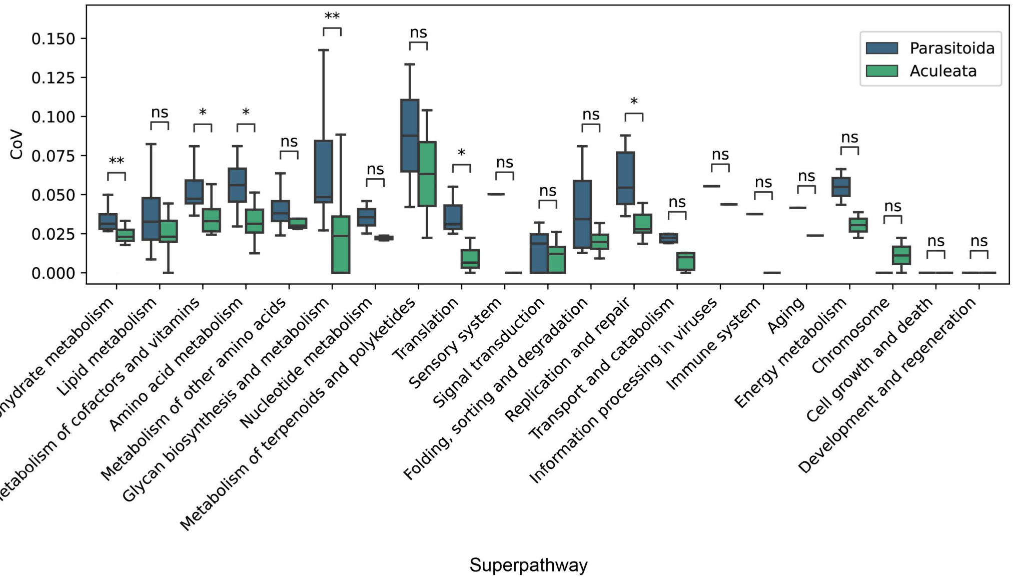


**Supplementary Fig. 18 | Comparison of the coefficient of variation (CoV) for pathway coverage in each superpathway in KEGG between Parasitoida and Aculeata.** The comparisons were performed using one-tailed Mann-Whitney *U* tests; *⩽05; **⩽0.01; *** ⩽0.001, ****⩽0.0001, and ns indicates no significant difference. The CoV of pathway coverage was calculated as the standard deviation divided by the mean of pathway coverage in species in Parasitoida and Aculeata, respectively.

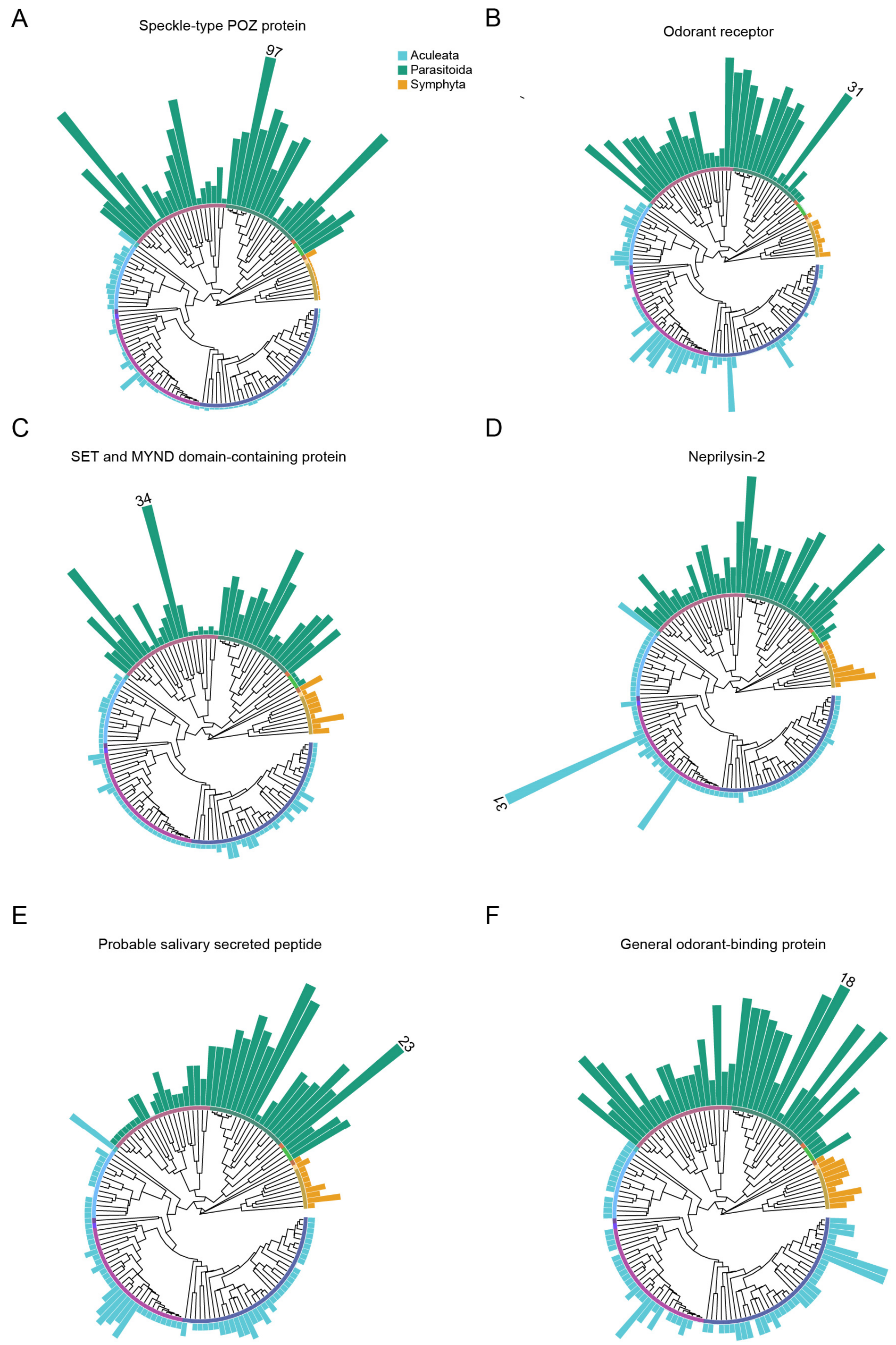


**Supplementary Fig. 19 | The size distribution of gene families significantly expanded in Parasitoida.** They were identified using one-sided Mann-Whitney *U* tests (FDR-adjusted $P < 0.05$). The number on the bar indicates the maximum gene copy count across the species for that family.

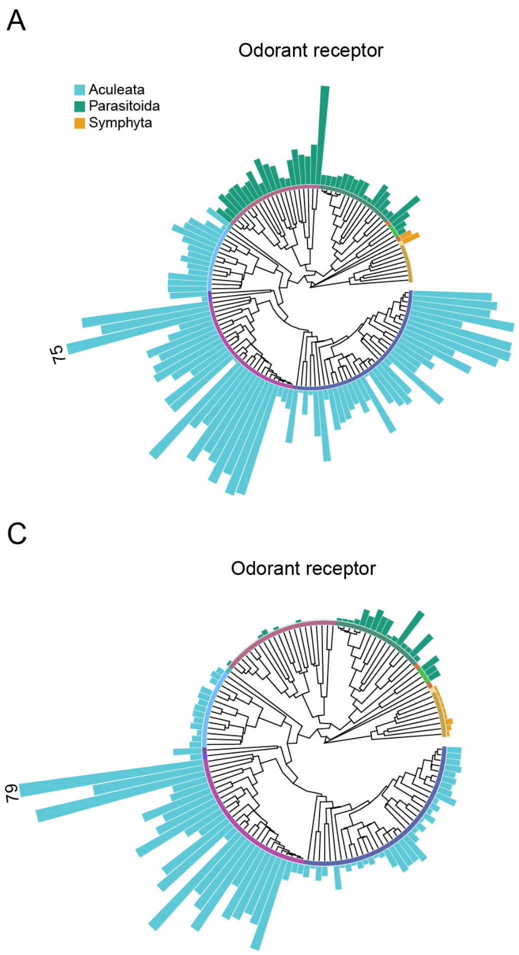


**Supplementary Fig. 20 | The size distribution of gene families significantly expanded in Aculeata.** They were identified using one-sided Mann-Whitney *U* tests (FDR-adjusted $P < 0.05$). The number on the bar indicates the maximum gene copy count across the species for that family.

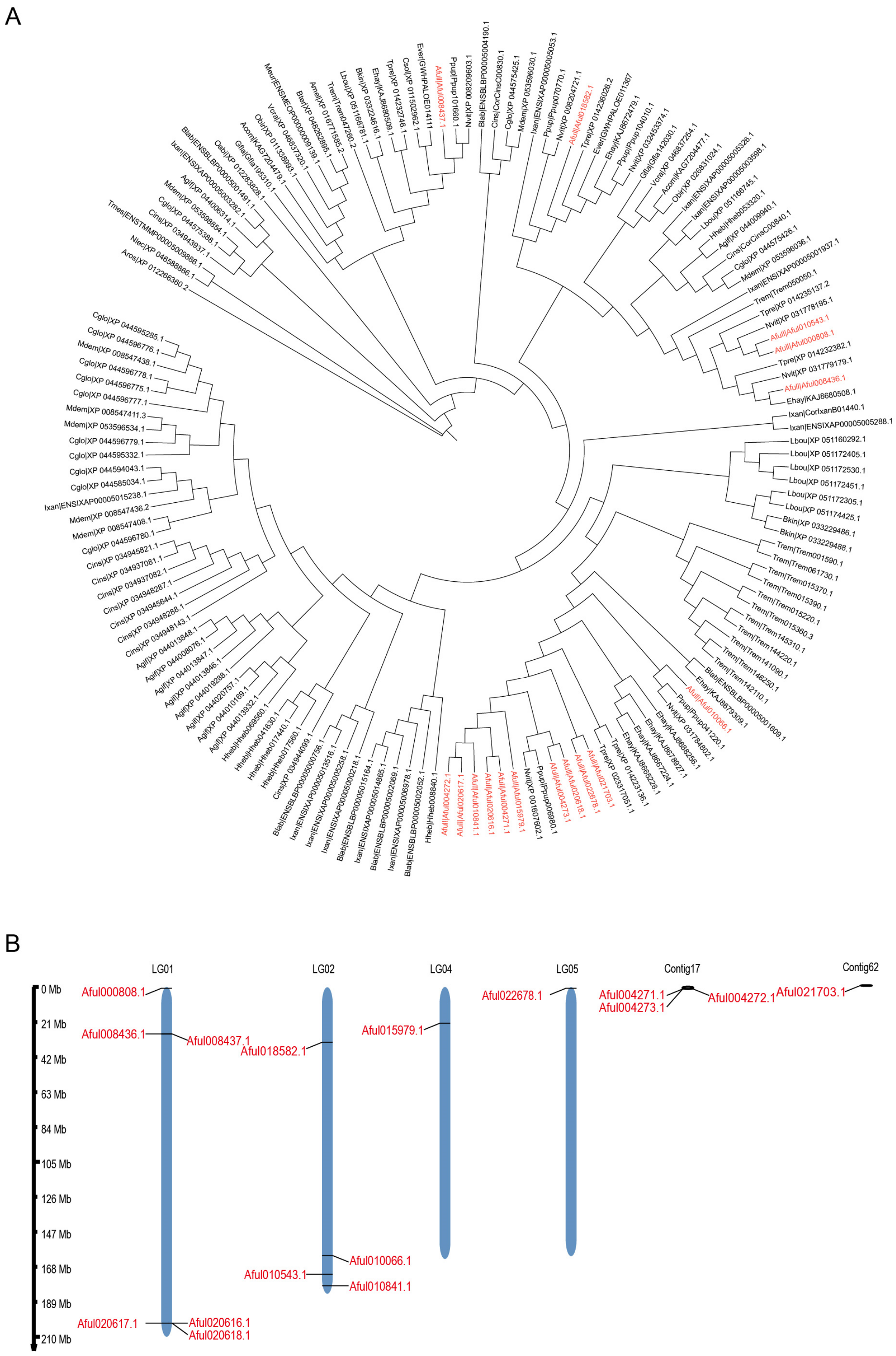


**Supplementary Fig. 21 | Evolutionary analysis of Venom metalloproteinase genes.** A) Phylogenetic analysis of Venom metalloproteinase genes in 28 hymenopteran species. The gene in red is for *Anastatus fulloi*. B) Distribution of Venom metalloproteinase genes on the *A. fulloi* genome.

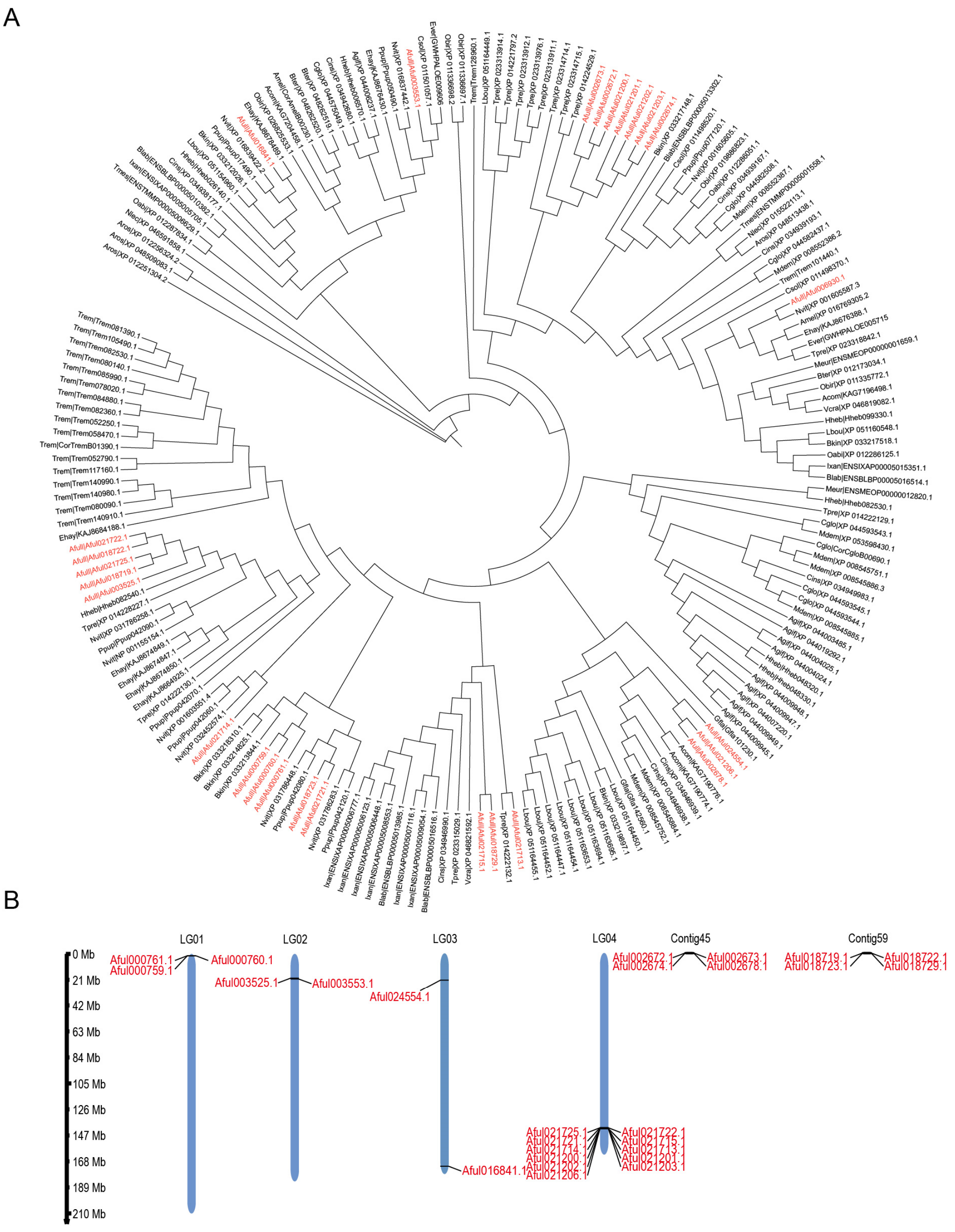


**Supplementary Fig. 22 | Evolutionary analysis of Venom allergen genes. A**) Phylogenetic analysis of Venom allergen genes in 28 hymenopteran species. The gene in red is for *A. fulloi*. B) Distribution of Venom allergen genes on the *A. fulloi* genome.

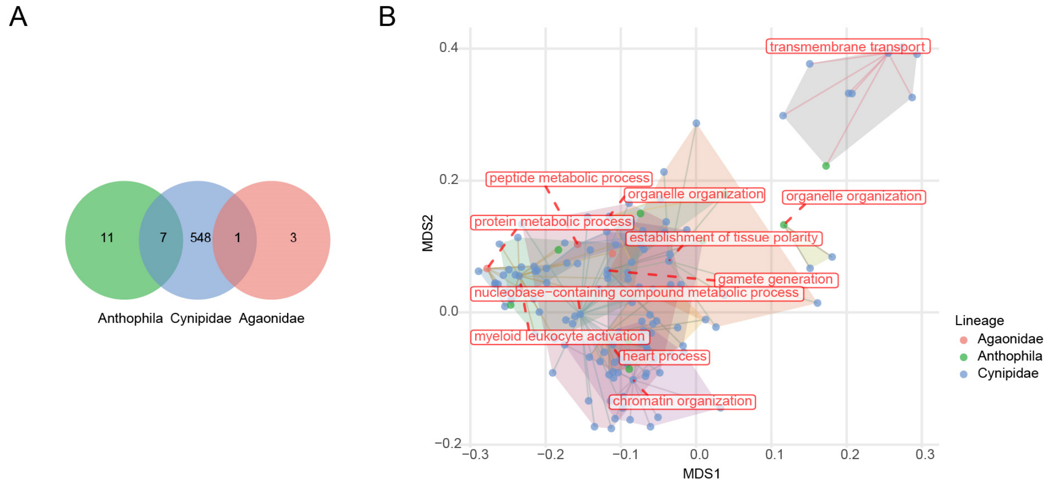


**Supplementary Fig. 23 | Functional convergence of exclusively expanded gene families in the three branches of secondary phytophagy origins.** A) The Venn diagram shows the number of shared and unique expanded gene families at three branches, with no expanded gene families shared by all three. B) The constellation plot shows the GO semantic space for the exclusively expanded gene families in the transitional ancestral branches. Each dot represents an exclusively expanded gene family in any of the three branches. The centroid gene family of each polygon is labeled with potential functions.

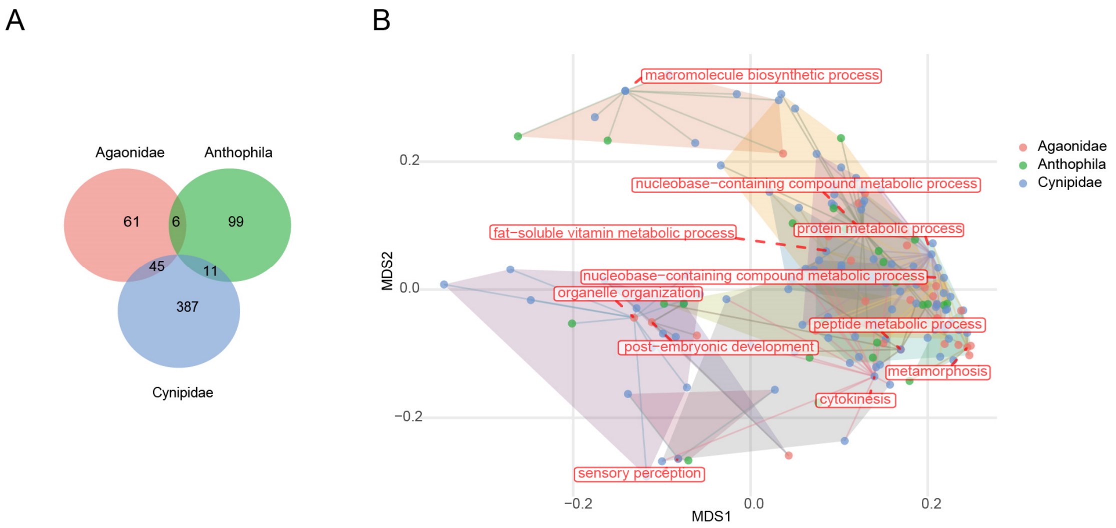


**Supplementary Fig. 24 | Functional convergence of exclusively contracted gene families in the three branches of secondary phytophagy origins.** A) The Venn diagram shows the number of shared and unique contracted gene families at three branches, with no contracted gene families shared by all three. B) The constellation plot shows the GO semantic space for the exclusively contracted gene families in the transitional ancestral branches. Each dot represents an exclusively contracted gene family in any of the three branches. The centroid gene family of each polygon is labeled with potential functions.

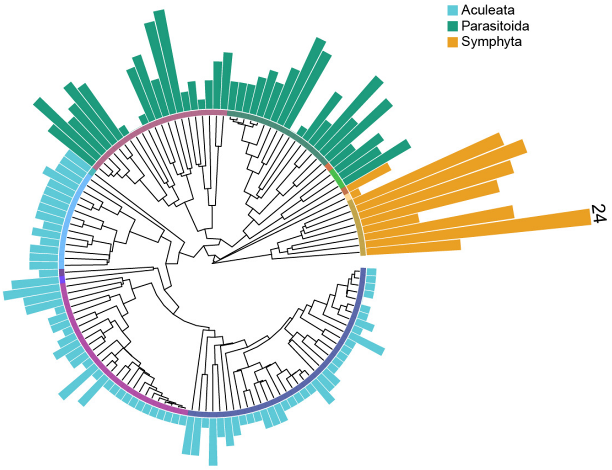


**Supplementary Fig. 25 | The size distribution of the myrosinase gene family in Hymenoptera.** The number on the bar indicates the maximum gene copy count across the species for the myrosinase gene family.
